# The telomere bouquet is a hub where meiotic double-strand breaks, synapsis, and stable homolog juxtaposition are coordinated in the zebrafish, *Danio rerio*

**DOI:** 10.1101/428086

**Authors:** Yana P. Blokhina, An D. Nguyen, Bruce W. Draper, Sean M. Burgess

**Affiliations:** Department of Molecular and Cellular Biology, University of California, Davis; Integrative Genetics and Genomics Graduate Group, University of California, Davis

## Abstract

Meiosis is a cellular program that generates haploid gametes for sexual reproduction. While chromosome events that contribute to reducing ploidy (homologous chromosome pairing, synapsis, and recombination) are well conserved, their execution varies across species and even between sexes of the same species. The telomere bouquet is a conserved feature of meiosis that was first described nearly a century ago, yet its role is still debated. Here we took advantage of the prominent telomere bouquet in zebrafish, *Danio rerio*, and super-resolution microscopy to show that axis morphogenesis, synapsis, and the formation of double-strand breaks (DSBs) all take place within the immediate vicinity of telomeres. We established a coherent timeline of events and tested the dependence of each event on the formation of Spo11-induced DSBs. First, we found that the axis protein Sycp3 loads adjacent to telomeres and extends inward, suggesting a specific feature common to all telomeres seeds the development of the axis. Second, we found that newly formed axes near telomeres engage in presynaptic co-alignment by a mechanism that depends on DSBs, even when stable juxtaposition of homologous chromosomes at interstitial regions is not yet evident. Third, we were surprised to discover that ~30% of telomeres in early prophase I engage in associations between two or more chromosome ends and these interactions decrease in later stages. Finally, while pairing and synapsis were disrupted in both *spo11* males and females, their reproductive phenotypes were starkly different; *spo11* mutant males failed to produce sperm while females produced offspring with severe developmental defects. Our results support zebrafish as an important vertebrate model for meiosis with implications for differences in fertility and genetically derived birth defects in males and females.

**Author Summary:** Inherent to reproduction is the transmission of genetic information from one generation to the next. In sexually reproducing organisms, each parent contributes an equal amount of genetic information, packaged in chromosomes, to the offspring. Diploid organisms, like humans, have two copies of every chromosome, while their haploid gametes (e.g. eggs and sperm) have only one. This reduction in ploidy depends on the segregation of chromosomes during meiosis, resulting in gametes with one copy of each chromosome. Missegregation of the chromosomes in the parents leads to abnormal chromosome numbers in the offspring, which is usually lethal or has detrimental developmental effects. While it has been known for over a century that homologous chromosomes pair and recombine to facilitate proper segregation, how homologs find their partners has remained elusive. A structure that has been central to the discussion of homolog pairing is the bouquet, or the dynamic clustering of telomeres during early stages of meiosis. Here we use zebrafish to show that the telomere bouquet is the site where key events leading to homologous chromosome pairing are coordinated. Furthermore, we show that deletion of *spo11*, a gene required for proper recombination in most studied organisms, resulted in very different effects in males and females where males were sterile while females produced deformed progeny.

## Introduction

Meiosis is a process that generates haploid gametes via one round of DNA replication and two rounds of chromosome segregation. Typically, homologous chromosomes (homologs) separate during meiosis I, and sister chromatids separate at meiosis II. Errors at either stage can lead to the production of aneuploid gametes, which is a major contributor to miscarriage and birth defects in humans [1]. During meiosis I prophase, homologs undergo pairing and crossing over, which is essential for their proper segregation in nearly every organism studied to date [2, 3]. Crossovers are created through a process termed recombination, where programmed DNA double-strand breaks (DSBs) are repaired using the homolog as a template, resulting in the exchange of homologous chromosome arms [2, 3]. Meiotic DSBs are formed by Spo11, a conserved topoisomerase-like enzyme [4]. They are then processed to reveal 3’ single stranded stretches of DNA that bind the strand exchange proteins, Dmc1 and Rad51, and engage in homology search [4–6]. In meiosis, the repair template is preferentially skewed towards the homologous chromosome rather than the sister chromatid, thus facilitating the pairing process [2]. While hundreds of DSBs might form in a single meiotic cell, only a subset go on to form crossovers, with the others being repaired via a noncrossover pathway [2, 3, 7].

DSB formation and crossing over occur in the context of the chromosome axis, a proteinaceous structure to which a linear array of chromosome loops is attached; both sister chromatids of a homolog are attached to a single axis and held together by cohesins [7, 8]. In many organisms, including mouse, budding yeast, and several plants, recombination initiation and repair intermediates are necessary for formation of the synaptonemal complex (SC), a tripartite proteinaceous structure that forms between the two homolog axes and holds them together along their lengths [2, 3, 7–9](Sup. Fig. 1). However, DSBs are dispensable for SC formation (synapsis) in some organisms such as *C. elegans, Drosophila*, and the planarian *Schmidtea mediterranea* [10–13]. In *C. elegans* and *Drosophila*, synapsis is initiated at “pairing centers” and ensues without the need for recombination [12]. In most organisms studied to date, recombination and the SC are used to link the homologous chromosomes. Notable exceptions are *Drosophila* males [14, 15] and *Lepidopteran* females [16] which do not form crossovers, and *Tetrahymena* [17] and fission yeast [18] which do not form the SC.

**Fig. 1.**
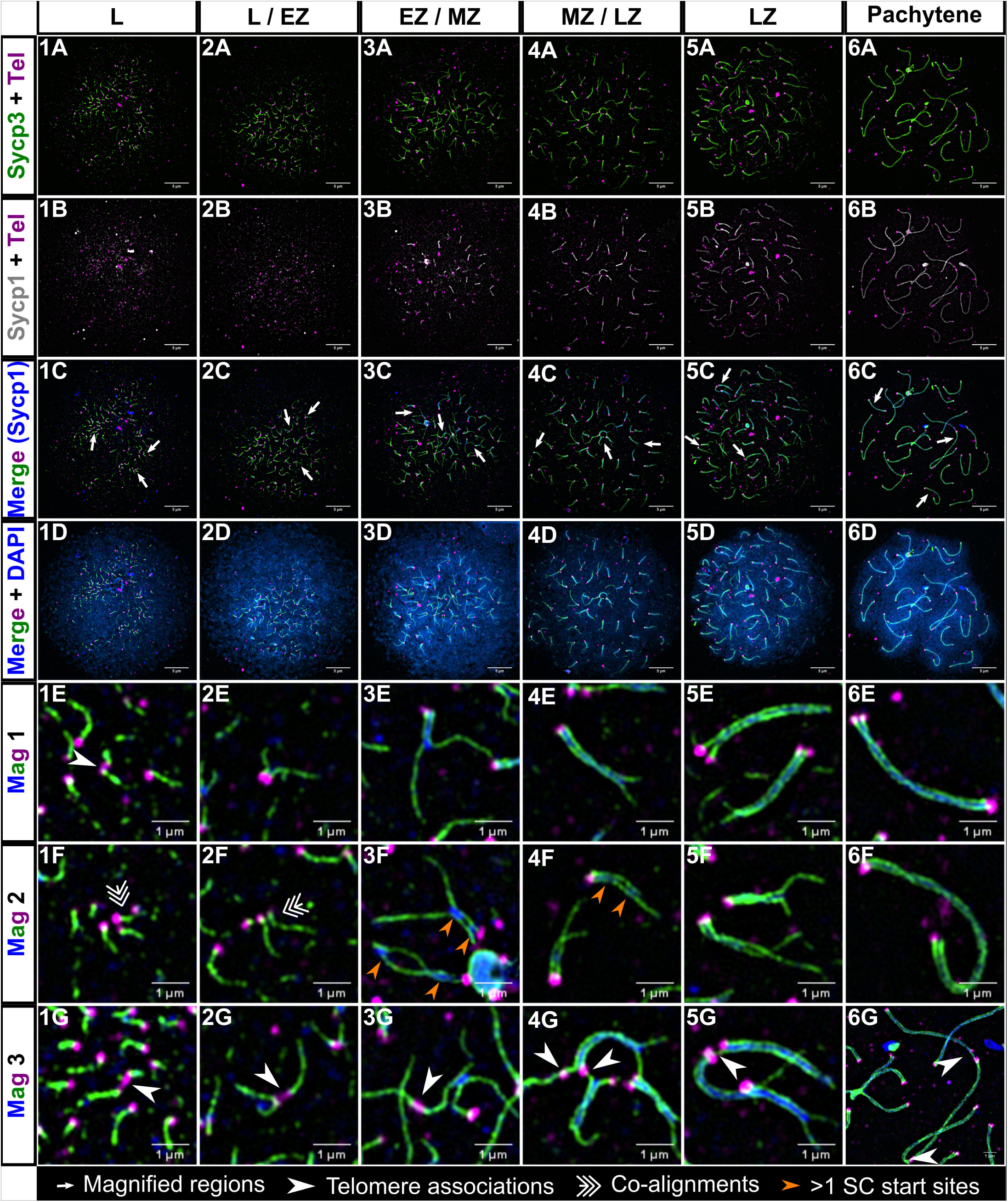
Homologous chromosome synapsis in male zebrafish meiosis I. 1A-6A: Loading of the chromosome axis protein, Sycp3 (green), initiates near the telomeres (Tel; magenta) in the bouquet and extends inward from both ends of the chromosome during meiotic prophase progression at the indicated stages (leptotene (L); leptotene to zygotene transition (L/EZ); early to mid-zygotene (EZ/MZ); mid to late zygotene (MZ/LZ); late zygotene (LZ); pachytene). 1B-6B: Sycp3 loading is closely followed by the initiation of SC formation, marked by Sycp1 (gray in row B to highlight features of the SC, blue in following rows)). 1C-6C: Merge of Sycp3, Sypc1 and Tel staining. 1D-6D: Merge of all signals plus DNA staining with DAPI. 1E-6G: Magnifications of three different regions (Mag 1, Mag 2, and Mag3) for each of the corresponding panels above; each magnified region is specified in images 1C-6C by long arrows. 1E, 1G-6G: Telomere associations are seen at all stages (arrowheads). 2G: Telomere association between three telomeres. 1F-2F: Short stretches of pre-synaptic co-alignment (triple arrows). 3F-4F: Chromosomes with more than one synapsis initiation site (orange arrows). 1A-6D scale bar = 5 μm. 1E-6G scale bar = 1 μm.

SC formation initiates primarily near telomeres in many organisms, including human males [19, 20], cattle males [21], the silkworm *Bombyx mori* [22, 23], the planarian *Schmidtea mediterranea* [13], and some plants such as tomato [24] and barley [25, 26]. In mouse males, while synapsis initiates interstitially as well as near the telomeres, there is a skew toward initiation at chromosome ends [27]. By contrast, synapsis in mouse and human females initiates primarily in interstitial regions [20, 28], while synapsis in female cattle initiates both near telomere ends and interstitially [21]. In many organisms, SC is nucleated preferentially at crossover fated sites [2]. Correspondingly, in mouse, human, and cattle, there is a skew toward crossovers in the distal regions of chromosomes in males but not in females [20, 29, 30].

During meiosis, telomeres are tethered to the nuclear envelope and their movement is directed by cellular cytoskeleton components [31–37]. One type of motion that is prominent in many species is the movement of chromosomes into and out of the bouquet, a conserved arrangement of chromosomes where telomeres are clustered together to one side of the nucleus. The bouquet has been hypothesized to restrict the chromosomes to one region of the nucleus thereby facilitating homolog recognition and pairing, possibly by limiting the homology search area or by active chromosome motion to disrupt weak non-specific interactions [2, 38]. However, in some organisms the bouquet does not exist (e.g. *C. elegans* and *Drosophila)* or does not form until after homologs are already co-aligned (budding yeast, *Sordaria*, mouse, and some plants) [2, 39], in which case it may play additional roles such as removing interlocks that form between two synapsed chromosomes [40]. Previous studies have shown that telomeres can form end-to-end associations in mammalian spermatids [41, 42] as well as in somatic and pachytene cells of some plants such as the dandelion-like smooth hawksbeard, *Crepis capillaris* [43, 44], pachytene cells of the cricket, *Gryllus argentinus* [45], and human spermatocytes [46]. The number and timing of these associations during meiotic prophase is poorly understood.

Our understanding of meiosis has been facilitated by the breadth of model organisms that have been studied, with each contributing new insight into the process. Budding and fission yeasts, *C. elegans, Drosophila*, mouse, and several plants have been instrumental to the study of the chromosome events of meiosis [9, 47]. While the basic features of meiosis are well conserved, the order of events and their functional dependencies vary significantly across species [2, 3, 48]. Indeed, the analysis of species-specific features has greatly informed our understanding of the seemingly fluid relationships between double-strand breaks (DSBs), the telomere bouquet, homolog pairing, synapsis, and recombination over evolutionary time scales [47]. There are remarkable similarities relating pairing, synapsis and recombination across phyla, even though difference in genome sizes can vary by orders of magnitude, especially when comparing mouse and humans to yeast, *Drosophila*, and *C. elegans*. Understanding how the chromosome events of meiosis are accommodated by larger genomes (and vice versa) necessitates the inclusion of additional model genetic organisms.

Zebrafish has many advantages as a vertebrate model for meiosis. Zebrafish can produce hundreds of offspring from a single cross, and external development allows for early detection of developmental abnormalities, including those caused by aneuploidy [49–51]. Unlike mammalian females, zebrafish females produce new oocytes throughout adulthood [52, 53], simplifying the characterization of female meiosis, which occurs in the fetal ovary in mammals. In addition, transparent gonads allow for observation of multiple stages of meiosis in a whole mount. Several cytological studies have provided insights into some aspects of zebrafish meiosis. For example, it has been shown in males that DSBs and the initial loading of the chromosome axis protein, Sycp3, and the SC transverse filament protein, Sycp1, are polarized to one side of the nucleus near the bouquet [6, 54–56]. Mlh1 foci, indicating sites of crossovers, have been shown to be distally skewed in males but not females [57]. These data are consistent with the genetic map where recombination is skewed to the telomeres in males yet more evenly distributed in females [58, 59]. A major gap of knowledge is that the order of events and the relationship between chromosome structure and recombination are not known. For example, it is not known if Spo11, a protein required for the formation of meiotic DSBs, is necessary for synapsis and homolog pairing in the zebrafish. This is an important relationship to determine as it sets the stage for further analyses of chromosome dynamics.

In this study we set out to establish the relationship between DSBs, synapsis initiation, and the establishment of close, stable homolog juxtaposition (which we refer to here as pairing) in the zebrafish males and females. We analyzed the progression of chromosome synapsis and pairing, telomere interactions, and double strand break localization at the super-resolution level. We created a knockout mutation in the *spo11* gene and found that both pairing and synapsis in the zebrafish are Spo11-dependent. We found dramatic sex-specific outcomes from disrupting Spo11: although synapsis and pairing defects were similar between *spo11* mutant males and females, males were completely sterile while females were able to produce offspring, though with severe developmental defects. Our results establish zebrafish as a tractable vertebrate model for understanding the chromosome events of meiosis I prophase from an evolutionary vantage and opening new lines of research with implications for human fertility and genetically derived birth defects.

## Results

### Sycp3 loading and synapsis are temporally and spatially coupled events in zebrafish spermatocytes

To better understand the relationship between the bouquet, Sycp3 loading, and synapsis in zebrafish, we set out to find when and where these events occur relative to one another in the prophase I nucleus. We stained spermatocyte nuclear spread preparations using a fluorescently tagged PNA probe by in situ hybridization to mark repeated telomere sequences, an Sycp3 antibody to mark the chromosome axis, and an Sycp1 antibody to mark the SC. The images were collected using structured illumination microscopy (Fig. 1, Sup. Fig. 1).

A general overview emerged. The telomere bouquet was prominent at the leptotene and zygotene stages (Fig. 1, panels 1D-5D). Early Sycp3 loading occurred adjacent to the telomere probe and extended toward the middle of the chromosomes as meiotic progression advanced. When the average Sycp3 length reached about 1 μm from the end of the chromosome, Sycp1 lines appeared close to the telomeres and then extended inward; as Sycp3 lines extended, Sycp1 lines closely followed, yet lagging somewhat behind (Fig. 2A). Interestingly, some chromosomes appeared to have more than one synapsis initiation site, albeit they were still in close proximity to the telomere (Fig. 1, panels 3F-4F).

**Fig. 2.**
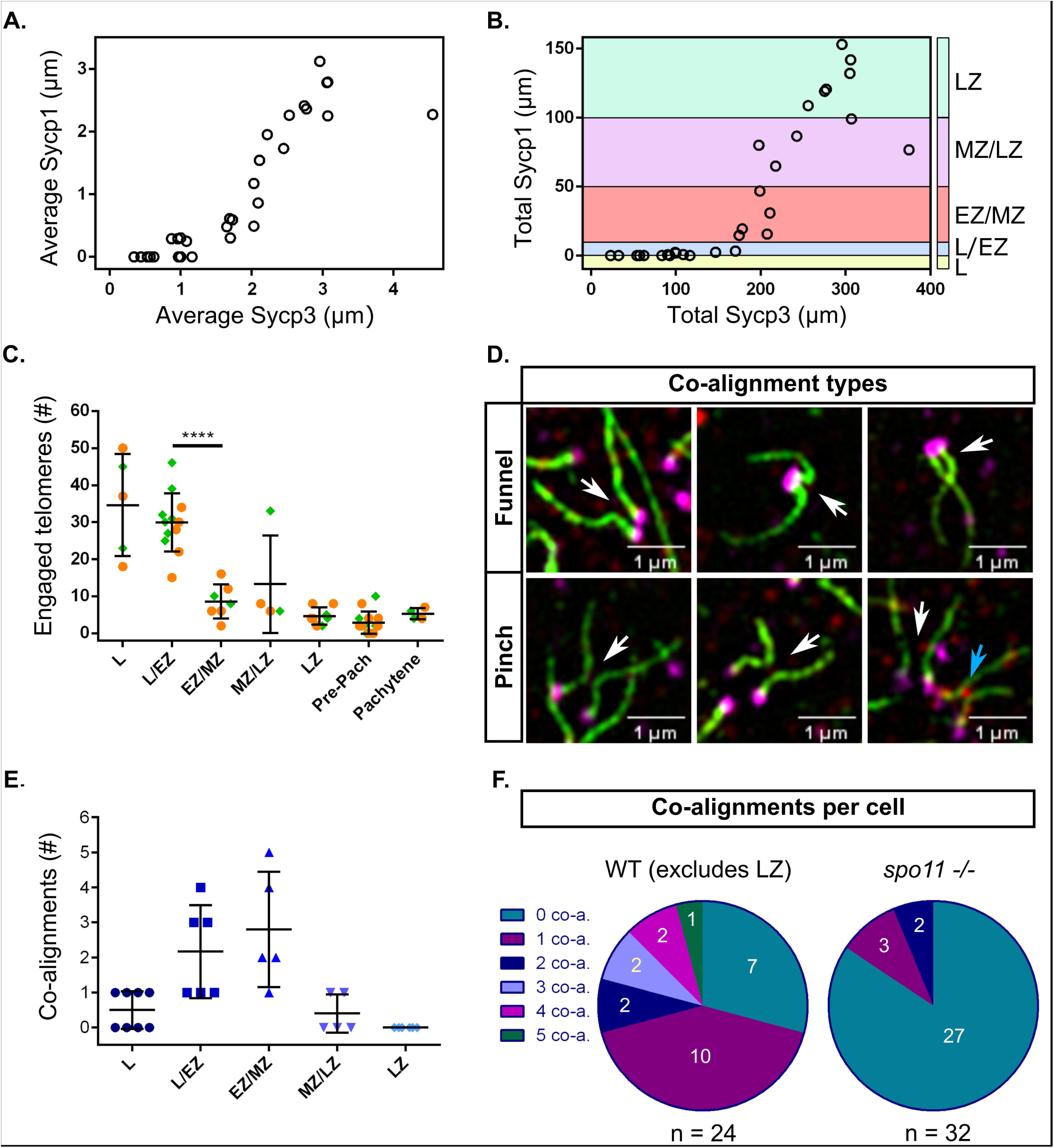
Measurements of Sycp3 vs. Sycp1 length, numbers of engaged telomeres, and numbers of co-alignments in male zebrafish meiosis I. A: Average Sycp3 length vs. average Sycp1 length. Each circle represents the average of all measurements in one nucleus (n = 30 nuclei). B: Total Sycp3 length vs. total Sycp1 length in the same cells shown in A. Stages of meiosis are defined by total length of Sycp1: leptotene (L, Sycp1 = 0 μm; yellow); leptotene to zygotene transition (L/EZ, Sycp1 = 1–10 μm; blue); early to mid-zygotene (EZ/MZ, Sycp1 = 10–50 μm; pink); mid to late zygotene (MZ/LZ, Sycp1 = 50–100 μm; purple); late zygotene (LZ, Sycp1 > 100 μm; cyan). C:Numbers of engaged telomeres per cell, where one engaged telomere end is an Sycp3 line with its telomere associated with a potentially non-homologous telomere end. Each point represents one cell. The data was pooled from two experiments done on different days (set 1, orange circles; set 2, green diamonds). The highest numbers of engaged telomere ends were observed in L and L/EZ stages after which point there was a significant decrease in engaged telomere ends in the EZ/MZ category, p < 0.0001 (unpaired t test with Welch’s correction), however, some number of engaged ends were maintained throughout prophase I. D: Examples of funnel and pinch co-alignment configurations. In the funnel configuration the co-alignment is directly adjacent to the telomeres; in the pinch configuration the co-alignment is not directly adjacent to the telomeres. White arrows point to the co-aligned regions. The blue arrow points to a pinch configuration where SC has already started to form (not counted as co-aligned). E: Total numbers of co-alignments in different stage cells of meiosis I prophase. Each point represents one cell. F: Numbers of co-alignments (0–5) in individual cells in WT (n = 24) and *spo11 -/-* (n = 32) males. The LZ class was excluded from the WT representation as at this stage synapsis has typically initiated at the majority of telomere ends. *spo11 -/-* cells, where synapsis is mostly abolished, were assessed for co-alignment if they appeared to have Sycp3 lines of similar lengths to the leptotene and zygotene classes of WT.

To facilitate comparisons between nuclei, we established staging criteria based on the total length of Sycp1 in 30 pre-pachytene images (Fig. 2B). We divided the nuclei into five classes: leptotene (L; Sycp1 = 0 μm), leptotene to early zygotene transition (L/EZ; Sycp1 = 1–10 μm, with few short stretches of SC near the telomeres), early to mid-zygotene (EZ/MZ; Sycp1 = 10–50 μm), mid- to late zygotene (MZ/LZ; Sycp1 = 50–100 μm), and late zygotene (LZ; Sycp1 > 100 μm). Spreads that contained more than three fully synapsed chromosomes were not measured as they were considered to be transitioning to pachytene and were referred to as “pre-pachytene”.

### Nonhomologous telomere associations are common throughout prophase I

In wild type zebrafish, we observed frequent end-to-end associations between telomeres, either as doublets or in higher order structures at all stages of prophase I (Fig. 1, panels 1E and 1G-6G, Fig. 2C). We assessed the extent of these associations by counting the total number of engaged telomere ends, defined by an Sycp3 line and its telomere associating with another end. One association could involve two or more engaged telomere ends. For a detailed description of our criteria see the methods. We observed the highest numbers of associations at the L to L/EZ stages, with an average of ~ 31% of telomeres engaged in associations with each other (Fig. 2C). The associations subsequently decreased to an average of ~ 6% but were still detected throughout the later stages including pachytene. In some cases, a short stretch of Sycp3 could be seen bridging the telomeres of unrelated chromosomes (Fig. 1, panels 5G-6G). The nature of these bridges is not known, however, Sycp3 protein can both bind dsDNA and form self-assemblies which are consistent with what we see [60, 61]. Associations involving more than two chromosomes indicated that at least a subset of telomere associations occurred between nonhomologous chromosomes or at the opposite end of the same chromosome. This was especially evident at the zygotene stage where telomeres of synapsing chromosomes were seen forming associations with a non-partner telomere, and at the pre-pachytene and pachytene stages where telomeres of synapsed chromosomes associated with nonhomologous chromosomes (Fig. 1, panel 6G). Our data suggest that telomere associations are a normal part of zebrafish meiosis I and not a pathological occurrence such as fusions caused by nonhomologous end-joining.

### Presynaptic co-alignment of homologous chromosomes occurs near the telomeres

In nearly every organism studied to date, axes of homologous chromosomes undergo some degree of co-alignment, or pairing, prior to synapsis, in which the distance between co-aligned axes is typically about 0.4 μm [2, 62]. Analysis of co-alignments seen in several species suggest the chromosomes are held in close proximity by DNA intermediates of the homologous recombination pathway [2]. We inspected images of spermatocyte nuclear spreads for evidence of axis co-alignment in the absence of detectable SC, marked by Sycp1. A detailed description of co-alignment assessment is provided in the methods. In brief, chromosome regions were considered co-aligned when Sycp3 lines were closely juxtaposed with the narrowest region between them at a distance of < 0.5 μm (Fig. 1, Panels 1F-2F, Fig. 2D). We found that presynaptic co-alignment of chromosomes occurred near the telomeres and adopted two main types of configurations: funnel and pinch. In the funnel configuration, co-alignment occurred directly adjacent to the telomeres to form the stem of the funnel while the other ends of the Sycp3 lines diverged away from the stem. The diverging lines could either be long or short (Fig. 1, panels 1F-2F; Fig. 2D), cross each other, or even fold backward toward the stem. In the pinch configuration, the narrowest region between the Sycp3 lines did not occur directly adjacent to the telomere, but a short distance away (Fig. 2D).

We found a total of 23 funnel configurations and 10 pinch configurations among 24 wild-type spermatocyte cells from the L to the MZ/LZ stage. During leptotene, co-alignments were rare, indicating that chromosomes were not stably juxtaposed with their partners at this stage (Fig. 2E). We found the highest number of co-alignments during the L/EZ and EZ/MZ stages, when the chromosomes begin to actively engage with each other to initiate synapsis. By the MZ/LZ and LZ stages, we found almost no co-alignments since most or all of the chromosomes had already engaged in telomere proximal synapsis (Fig. 2E). As individual nuclei with presynaptic co-alignment usually have less than a total of five co-alignments (Fig. 2F), we believe that this is a transient stage mediated by homologous recombination that quickly progresses to synapsis initiation for any given chromosome pair. The funnel and pinch configurations likely represent recombination events that initiate very close to the telomere or slightly inward. It is likely that the pinch and funnel axis shapes are precursors to synapsis initiation since we saw similar configurations with short stretches of SC (Fig. 1, panels 2E and 2F; Fig. 2D). Moreover, unpaired telomeres in the pinch configurations suggest that stable telomere associations are not the primary driver of homolog pairing and that initial homology recognition can occur in sub-telomeric regions.

### Co-alignment at interstitial loci is coincident with synapsis

While we observed pre-synaptic co-alignment near the telomeres, we found no evidence for similar co-alignment at interstitial regions of the axes. For chromosomes where synapsis had initiated, the distances between the diverging edges of the two Sycp3 lines were often much greater than 0.4 μm and were frequently bent in non-parallel orientations with respect to each other (Fig. 1, panels 2E, 3E, 3F, 5F, 3G, 4G). This suggested that axes were not stably co-aligned at interstitial regions prior to synapsis. To determine when interstitial sites become stably co-aligned relative to SC formation, we performed fluorescence in situ hybridization (FISH) on cells at different stages of meiotic prophase I using a 68 kilobase bacterial artificial chromosome (BAC) probe located ~10 Mb from the end of chromosome V (total 72.5 Mb) (Fig. 3, Sup. Fig. 2). We found that the BAC signals were far apart in the early prophase I stages but paired up as synapsis progressed (EZ/MZ to LZ). In a few instances, we found cells where the BAC probe localized to forked regions of Sycp3 just ahead of synapsis (Fig. 3C). These data suggest that stable juxtaposition between homologs does not occur until they are synapsed at that region.

**Fig. 3.**
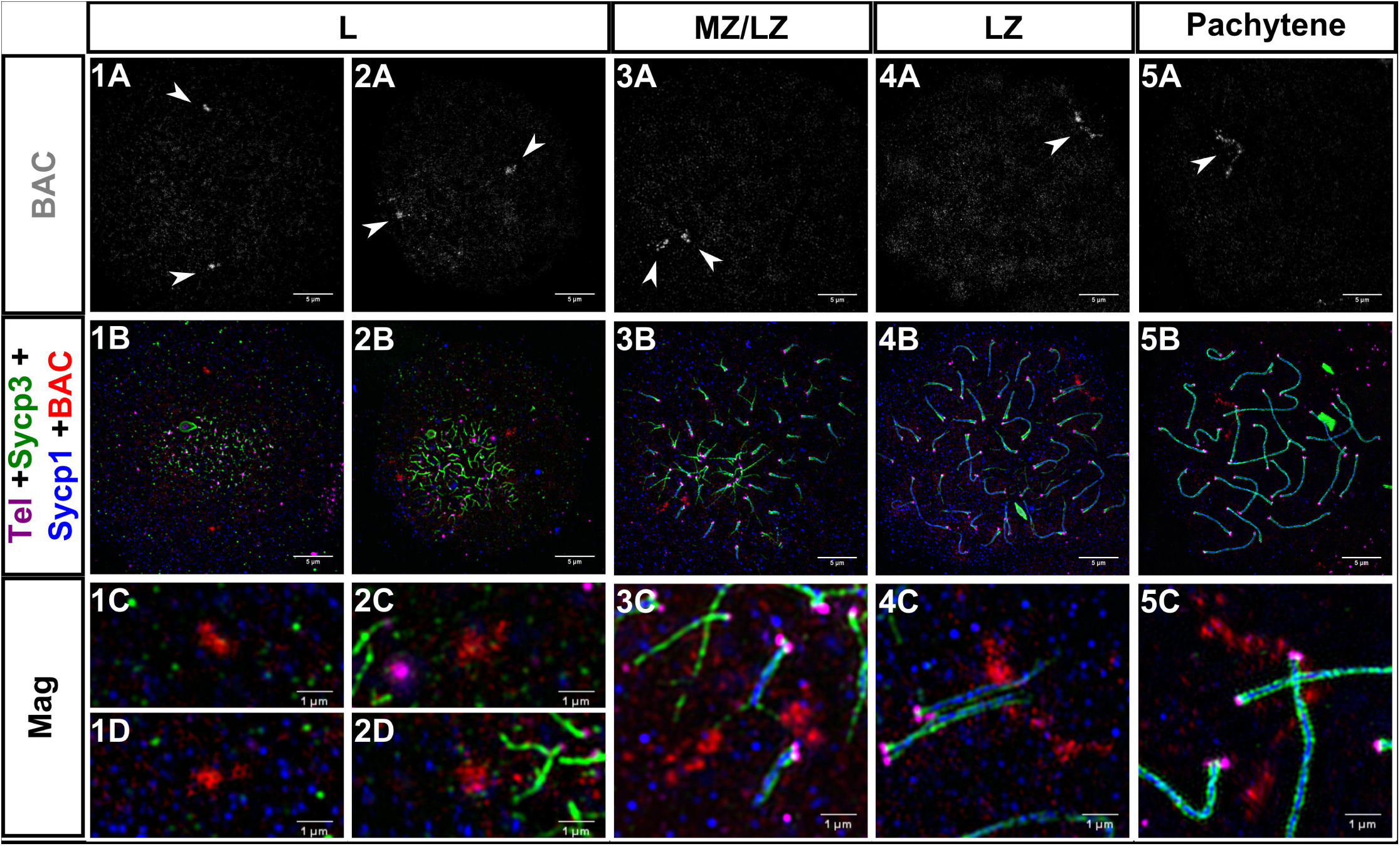
Homologous chromosome pairing at interstitial sites is coincident with synapsis in zebrafish male meiosis I. 1A-5A: Fluorescent in situ hybridization with a BAC probe (gray, white arrowheads) to a region ~10Mb from the telomere on chromosome 5. 1B-5B: Same images as 1A-5A plus telomeres (Tel; magenta), Sycp3 (green), and Sycp1 (blue); the BAC is in red. 1C-5C, 1D-2D: Magnifications (Mag) for each of the corresponding panels above. 1A-5B scale bar = 5 μm. Mag scale bar = 1 μm. Leptotene (L), mid to late zygotene (MZ/LZ); late zygotene (LZ). Note that images for panel series 3 and 5 are from a spreading procedure performed on a separate day from panel series 1, 2, and 4, and are included to illustrate a rare observation where BAC probes are located on axes where SC is extending (3C) and also where the BAC signal is highly elongated (5C).

Prior to pairing, the BAC signal presented as an amorphous shape, while at synapsed regions the shape was elongated and appeared to lie perpendicular to the axis (Fig. 3, Sup. Fig. 2, LZ and Pachytene panels). The two prominent features of a meiotic chromosome are the axes that run along the chromosome length and the DNA loops that attach to the axis. It is possible that the change in shape of the BAC signal reflects a change in chromosome architecture, for example, from a fractal globule, characteristic of interphase chromosomes [63], to a looped region that characterizes the DNA component of meiotic chromosomes [62].

### Females progress through prophase I similarly to males

Several lines of evidence point to differences between female and male meiosis in zebrafish. Females have longer chromosome axes and exhibit a more even distribution and higher numbers of crossovers [57–59, 64]. In order to determine if early prophase events in females differed from males, we stained ovary nuclear surface spreads with the telomere probe and antibodies against Sycp3 and Sycp1. We staged nuclei based on the overall appearance of axis extension and synapsis progression. We found that the progression of prophase I in females was similar to that in males: Sycp3 loading and synapsis initiated near both telomere ends in the bouquet and elongated toward the center of the chromosome, with synapsis lagging behind Sycp3; chromosomes appeared to become stably juxtaposed as synapsis progressed (Fig. 4).

**Fig. 4.**
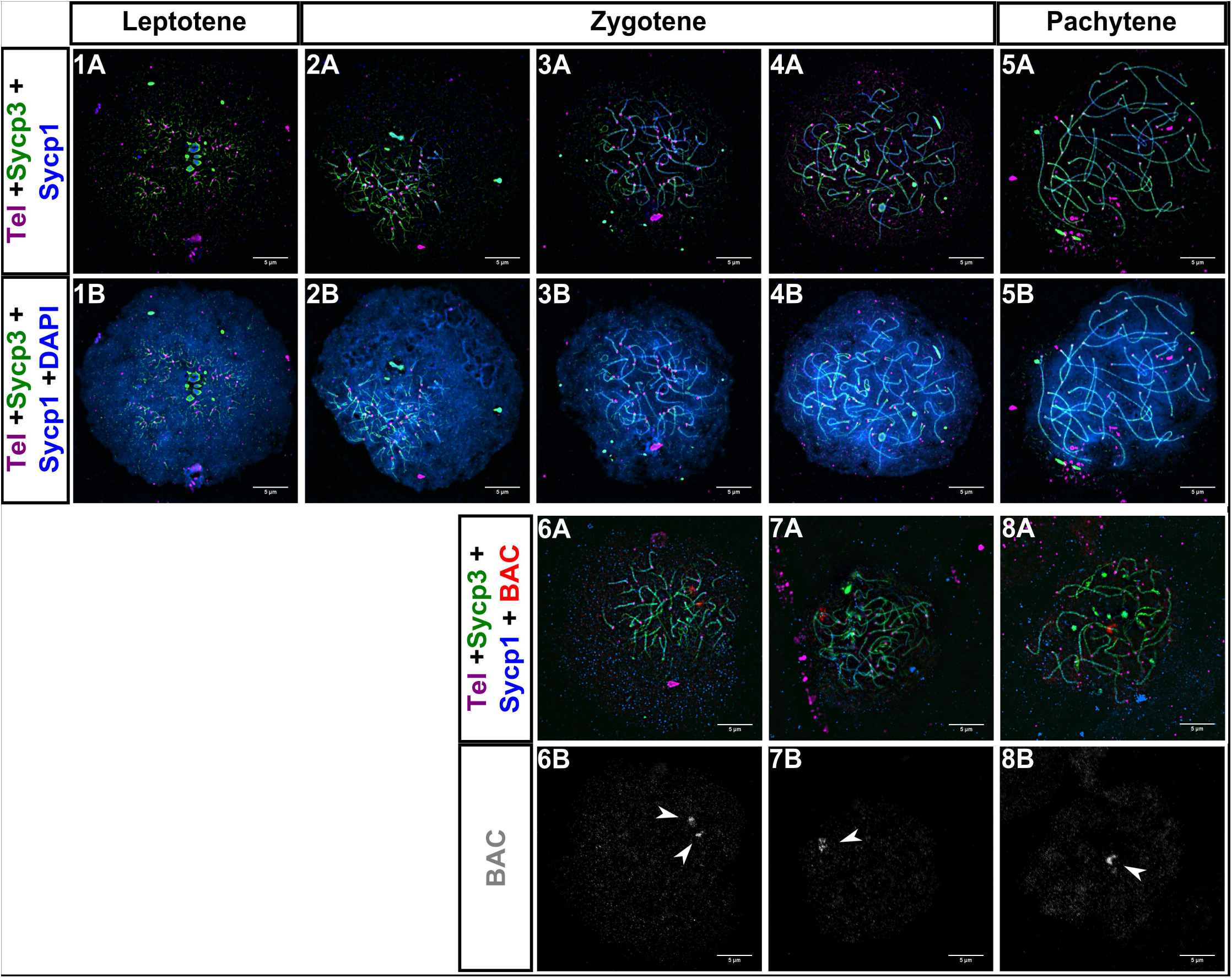
Homologous chromosome synapsis and pairing in female zebrafish meiosis I. 1A-5B: Sycp3 (green) loading and synapsis (Sycp1; blue), begin near telomeres (Tel; magenta) and progress inwards, similarly to what is seen in zebrafish males. 6A-8A: BAC (red), Sycp3 (green), Sycp1 (blue), Telomeres (Tel; magenta). 6B-8B: Same images as 6A-8A with only the BAC shown (gray; arrowheads). Scale bar = 5 μm.

### Chromosome interlocks and regional de-synapsis are common during the late zygotene to pachytene transition

Chromosome interlocks are common during the zygotene stage in several organisms including the silkworm, *Bombyx mori*, and maize, *Zea mays*, but most interlocks are resolved by pachytene [22, 62, 65, 66]. In the zebrafish spermatocytes, we regularly found chromosomes that were intertwined or sometimes trapped between another set of homologs at the pre-pachytene stages when most of the chromosomes were already synapsed (i.e. 16 - 24 fully synapsed chromosomes; Fig. 5). These configurations closely approximated interlock structures seen at late zygotene in other species [2]. Interlocks occur when one chromosome or a pair of chromosomes becomes entrapped between the space of two synapsing homologs. Thus, from a first approximation, the structures we see are likely interlocks. Interestingly, nuclei at this stage frequently also had individual pairs of chromosomes with extensive or complete de-synapsis, sometimes with another homolog appearing to be entrapped in the desynapsed region (Fig. 5, Panels 1A-1C, 2A-3D, 5A-6D). Of 8 cells at this pre-pachytene stage, only 2 showed no anomalies. Of the remaining cells, 3 had both interlocks and de-synapsis, 1 had just interlocks, and 2 had just de-synapsis. The de-synapsis is unlikely to be due to the cells transitioning to diplotene, as some de-synapsed chromosomes were completely separated with no evident crossover connections, and some were entangled around other chromosomes. Although interlocks are not a common feature of pachytene cells, it is also possible that there is a subset of cells where interlocks persist through pachytene and the chromosome-wide de-synapsis we see are chromosomes in diplotene. Interestingly though, we never saw a spread nucleus showing a classic diplotene state as seen in other organisms where a complete set of de- synapsed bivalents were held together by one or more chiasmata, suggesting this state in males is transient or full-length Sycp3 axes start to degrade at this stage.

**Fig. 5.**
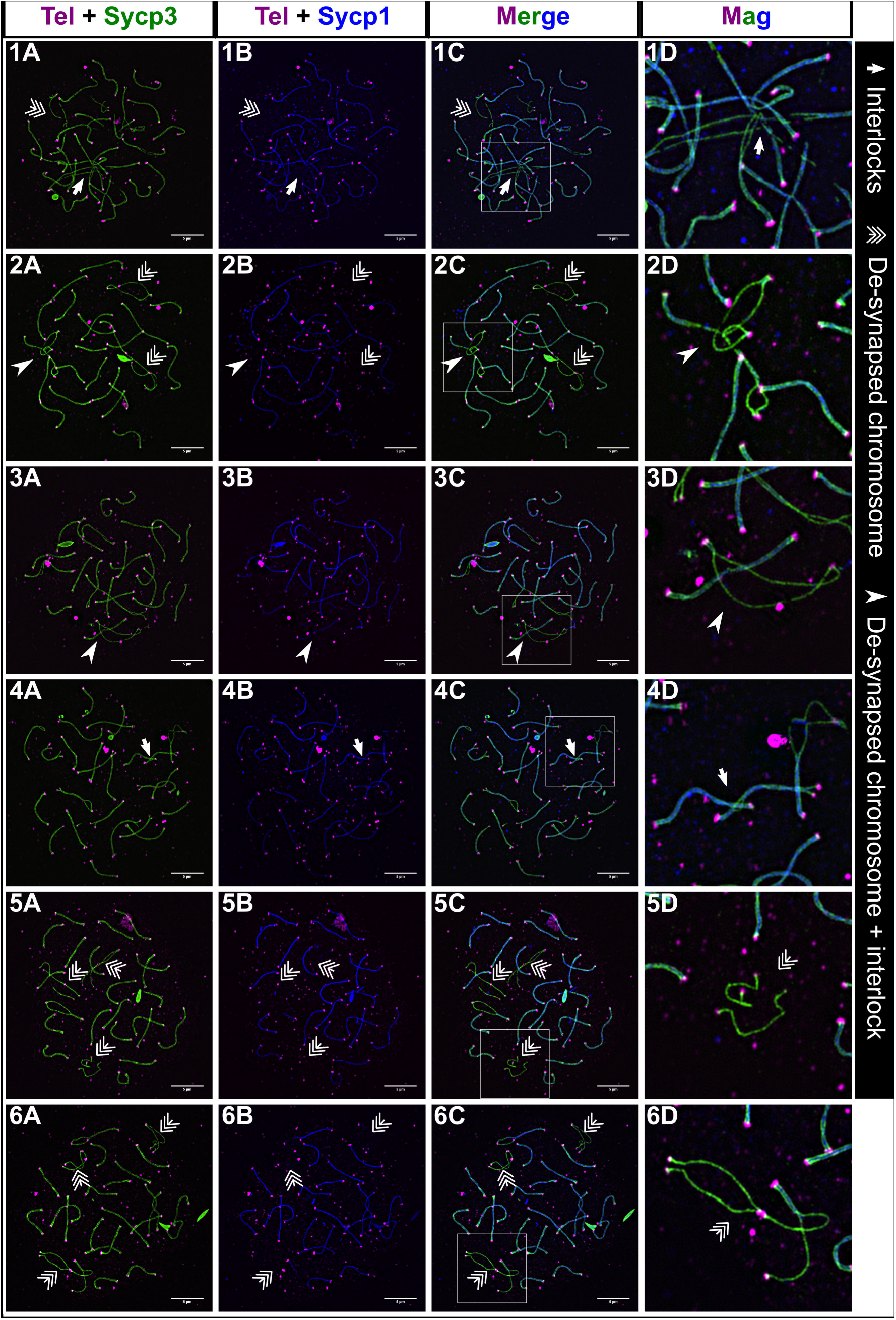
Interlocks and de-synapsed chromosomes in male zebrafish meiosis I. Examples of nuclear surface spreads showing interlocks (arrows) and de-synapsed chromosomes (triple arrows) at the late zygotene to pachytene stage where most chromosomes are synapsed. In some cases, de-synapsed chromosomes have another chromosome passing through them (arrowheads). Magnification (Mag) images are from regions indicated by white boxes in the merge panels. Sycp3 (green), Sycp1 (blue), Telomeres (Tel; magenta). Scale bar = 5μm.

### DSBs cluster near the telomeres in bouquet formation

Since the sites of crossovers in zebrafish are skewed toward the ends of chromosomes in males [57], we suspected that co-alignment and synapsis near telomeres might be initiated by local DSBs. We first tested if γH2AX, a biomarker for DSBs, co-localizes with telomeres in sectioned testes and found a sharp polarization of γH2AX staining to one side of the nucleus when chromosomes were in the bouquet and then a more dispersed signal in cells where chromosomes had exited the bouquet (Fig. 7A). These results are consistent with a study that showed γH2AX signal clustered with initiation of axis formation marked by Sycp3 [55].

To evaluate the distribution of DSBs at the super-resolution level, we probed spermatocyte nuclear surface spreads with telomere probes and antibodies to Sycp1 and the DSB repair protein Rad51. Previous work at lower resolution showed that Rad51 foci were primarily found near sites where Sycp3 loading had initiated [6]. Consistent with this finding, and with the γH2AX localization, we found that Rad51 foci were interspersed with the telomere foci in the bouquet cluster (Fig. 6B). If DSBs are required for initiating synapsis, then we expected to find that most SC stretches would be associated with a Rad51 focus. This was not the case, however, since there were many instances of SC stretches with no associated Rad51 foci (Fig. 6B, panels 1C-3C). We found that 39% (n = 269) of synapsed ends in early zygotene had no associated Rad51 focus. This was surprising given that we also found that synapsis requires Spo11-dependent DSBs (below). Three possible reasons could account for this observation: 1) some synapsis may occur independent of DSBs, 2) synapsis is initiated at Rad51-associated DSBs but Rad51 signal has been lost due to repair prior to imaging, or 3) some synapsis is initiated at Dmc1-associated DSBs that do not co-localize with the Rad51- associated DSBs. The latter is supported by data from *Arabidopsis thaliana* where Dmc1 and Rad51 DSBs do not colocalize [67].

**Fig. 6.**
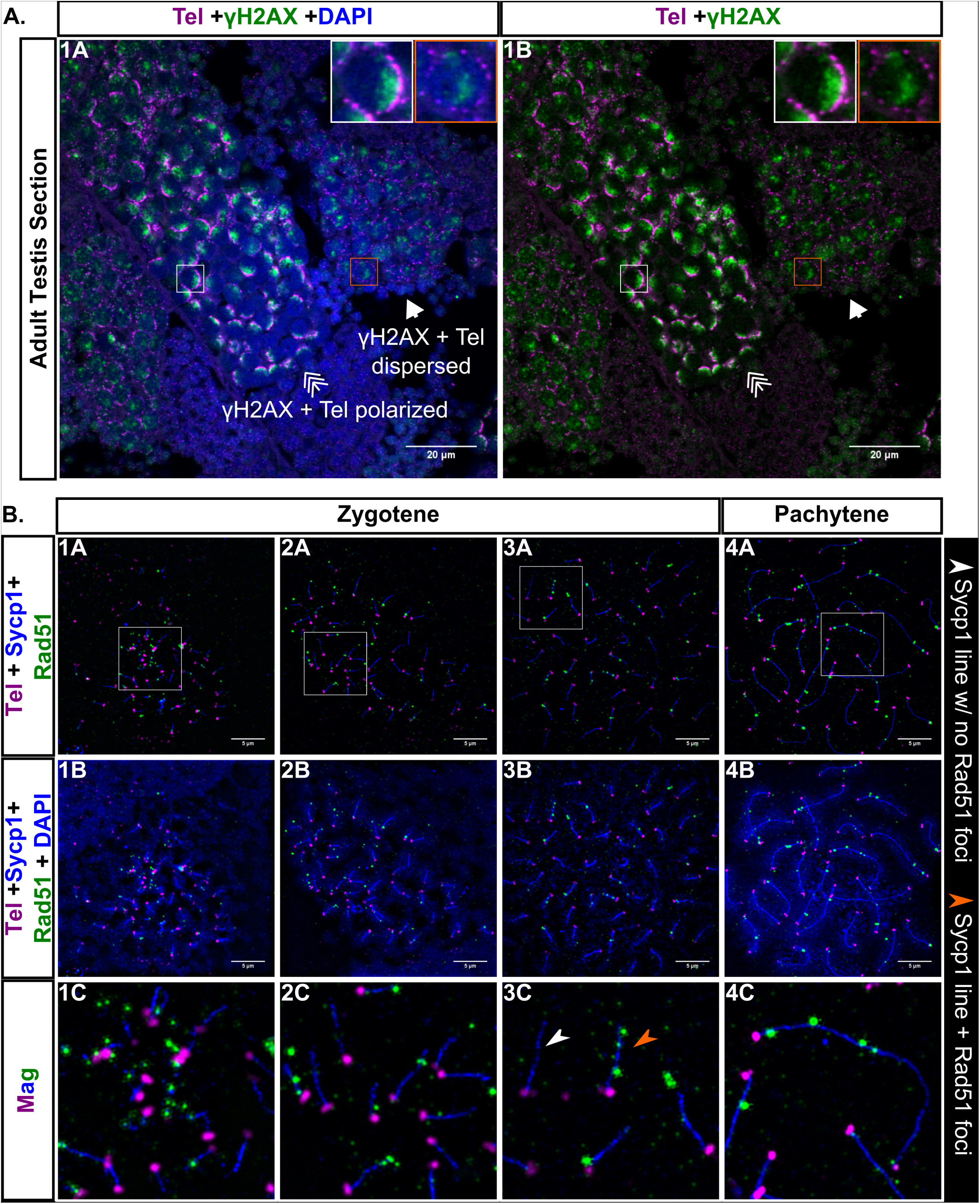
Double strand break (DSB) localization during male zebrafish meiosis I. A: Testis section stained for γH2AX (green), telomeres (Tel; magenta), and DNA (DAPI, blue). γH2AX staining is polarized when the telomeres are in bouquet formation (triple arrow, white boxes) and is more scattered when the telomeres are dispersed (arrow, orange boxes). Scale bar = 20 μm. B: Panels 1A-4C: Nuclear surface spreads stained for Sycp1 (blue), Rad51 (green), telomeres (Tel, magenta) and DNA (DAPI, blue); 1C-4C: Magnified (Mag) images are from regions indicated by white boxes in the top row panels. White arrowhead points to an example of an Sycp1 stretch with no Rad51 foci. Orange arrowhead points to an example of an Sycp1 stretch with Rad51 foci. 1A-4B scale bar = 5μm.

We found that some cells at pachytene had Rad51 foci and they were located both near the telomeres and in interstitial regions. There are two ways we can envision the interstitial foci could arise: 1) All meiotic DSBs form at the same time when the cells are in the bouquet stage, in which case DSBs at interstitial locations would be recruited to the bouquet, or 2) breaks continue to form throughout prophase I. Further studies are required to distinguish between these models. Combined, our results show that DSBs are primarily clustered near the telomeres but are also found at interstitial regions during pachytene, which reflects the crossover pattern in zebrafish males [57, 58].

### Synapsis and stable homolog juxtaposition, but not telomere associations or the bouquet, are disrupted in *spo11* mutant males and females

In order to determine whether synapsis can occur in the absence of Spo11, which is required for the formation of meiotic DSBs, we created a *spo11^−/−^* mutant. The *spo11* gene in zebrafish consists of 13 exons encoding a 383-amino acid protein product (GenBank: AAI65825.1) with the predicted TP6A_N superfamily domain at 96–157 aa and the predicted TOPRIM superfamily domain at 205–367 aa (NCBI BLASTP 2.8.0). We used TALENs targeted to the second exon to introduce an indel mutation by error prone repair. Sequencing of genomic DNA isolated from offspring of founder backcrosses identified an 11 bp deletion resulting in a frameshift mutation in the coding region that predicts a truncated protein of 57 aa lacking both the TP6A_N and TOPRIM domains (Fig. 7A). To confirm disruption of Spo11 function in the mutant, we probed whole mount testes of *spo11^−/−^* males with antibodies to γH2AX and the germ-cell specific Vasa protein and found that γH2AX clusters were absent in the germ cells, showing that the mutant is deficient for DSB formation (Fig. 7B).

**Fig. 7.**
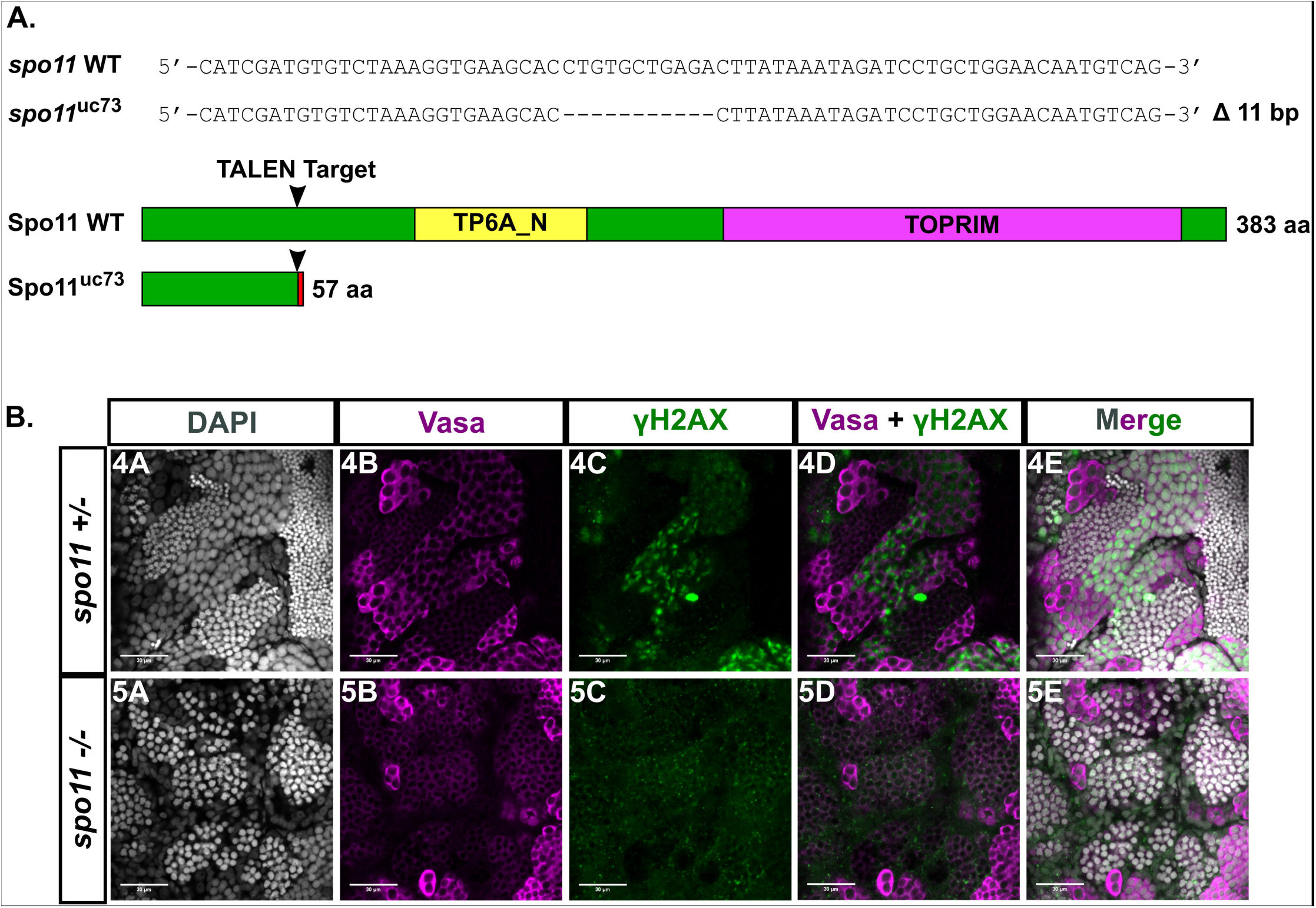
A: TALEN generated 11 bp frameshift mutation allele (*spo11*^uc73^) leads to a truncated 57 aa Spo11 protein with the conserved TP6A_N and TOPRIM family domains deleted. B: Whole testes stained for vasa (magenta), γH2AX (green), and DAPI (gray). *spo11 +/−* testes show a normal pattern of γH2AX staining whereas the pattern is disrupted in the *spo11 -/-* mutant.

We next examined evidence of synapsis and pairing in nuclear surface spreads from *spo11* mutants. Although Sycp3 loading initiated near the telomeres and elongated inward as in wild type, Sycp1 loading did not follow (Fig. 8A). We divided *spo11* mutant spermatocytes into the L - L/EZ-like, EZ-LZ-like, and Post-LZ-like categories based on the overall resemblance of Sycp3 loading in the nucleus to equivalent wild-type stages. Post-LZ included pre-pachytene-like or pachytene-like stages. In *spo11* mutant spermatocytes, 30 out of 40 cells had no synapsis and the remaining cells had between 1 and 4 short fragments of Sycp1, which appeared either between two axes, on one axis, or as a lone filament (Fig. 8A, panels 3C-4C). These Sycp1 stretches may have been due to self-assembly of Sycp1 filaments [68]. In addition, the bouquet was also maintained. Telomere-proximal co-alignment between chromosomes, however, was disrupted (Fig. 2F).

**Fig. 8.**
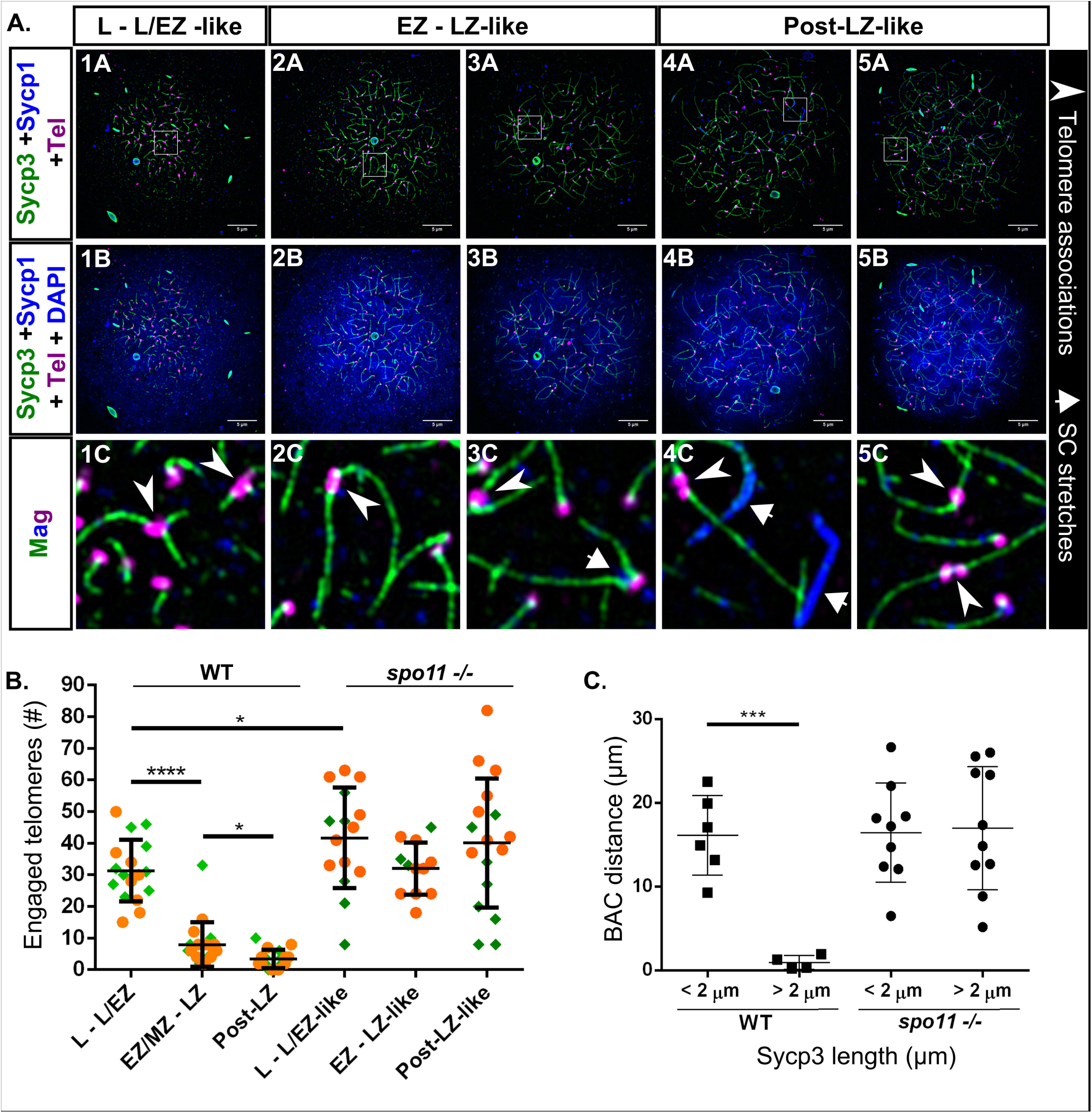
*spo11 -/-* males fail to properly synapse homologous chromosomes, and exhibit high numbers of engaged telomeres during meiosis I. A: 1A-5B: *spo11 -/-* nuclear surface spreads stained for Sycp3 (green), Sycp1 (blue), telomeres (Tel; magenta), and DNA (DAPI, blue). 1C-5C: Magnified (Mag) images are from regions indicated by white boxes in the panels above. Telomere associations are seen at all stages (arrowheads); Short stretches of Sycp1 are seen in some mutant cells (short arrows). Scale bar = 5μm. B: Numbers of engaged telomeres in WT vs. *spo11 -/-* cells. The mutant cells were categorized based on similarity to WT Sycp3 loading extent. Post-LZ refers to all cells that were classified as pre-pachytene or pachytene in WT and equivalent cells in *spo11* mutants. The data was pooled from two experiments done on different days (set 1, orange circles; set 2, green diamonds). P values were acquired using unpaired t test with Welch’s correction. For * p ≤ 0.0314, and for **** p < 0.0001. While WT telomere engagements are reduced at later stages, the telomere engagements in *spo11-/-* are maintained. C: Dot plot of BAC distance measurements in WT and *spo11 -/-* males (in μm). In the WT during the early stages when the average Sycp3 lines are short (< 2 μm), the BAC signals are unpaired; As meiosis progresses and the Sycp3 lines elongate (> 2 μm), the BAC signals exhibit pairing (p = 0.0004, unpaired t test with Welch’s correction). In the *spo11 -/-* males the BAC signals remain unpaired even with extended Sycp3 axes.

Intriguingly, we found an average of ~ 42% of telomeres engaged in associations in the L – L/EZ-like mutant cells as compared to the ~ 31% we see in equivalent stages of wild type (Fig. 8A, panels 1C-5C, Fig. 8B; p = 0.0389). Unlike in wild type, the telomere associations were maintained at high levels in the mutant throughout prophase I. Several possibilities could account for the loss of telomere associations in wild-type cells. Pairing and synapsis between the ends of homologous chromosomes could physically displace weak associations between non-related chromosome ends. Alternatively, a regulatory feature associated with the transition from leptotene to zygotene could signal loss of a subset of associations, or the reduction of entanglements later in meiosis could allow associations to be disrupted by the physical force of spreading. Any one of these possibilities could account for the persistence of associations in the *spo11* mutant.

Wild-type cells that were in the EZ/MZ to LZ stages gave a distribution of inter-BAC distances that were overall shorter than those in the L or L/EZ stage (0.24–3.13 μm vs. 9.3–25.3 μm, p = 0.0004, Fig. 8C). In the L to L/EZ stages the average length of Sycp3 lines was less than 2 μm while in the later stages, the average length of Sycp3 lines was greater than 2 μm. The sharp decrease in inter-BAC probe distance measurements suggests that pairing at the probed locus occurs shortly after synapsis is initiated. For the *spo11* mutant, we staged the nuclei based on Sycp3 axis length since the SC was absent. BAC foci in the *spo11* mutants remained at approximately the same distance from each other when the axes were short (< 2 μm) or long (> 2 μm) (Fig. 8C, Sup. Fig. 3). In mutant females, synapsis and pairing were also disrupted as was seen in males (Fig. 9, Sup. Fig. 3). Together, these data indicate that Spo11 is required for the initiation and/or stabilization of synapsis and homolog juxtaposition in males and females.

**Fig. 9.**
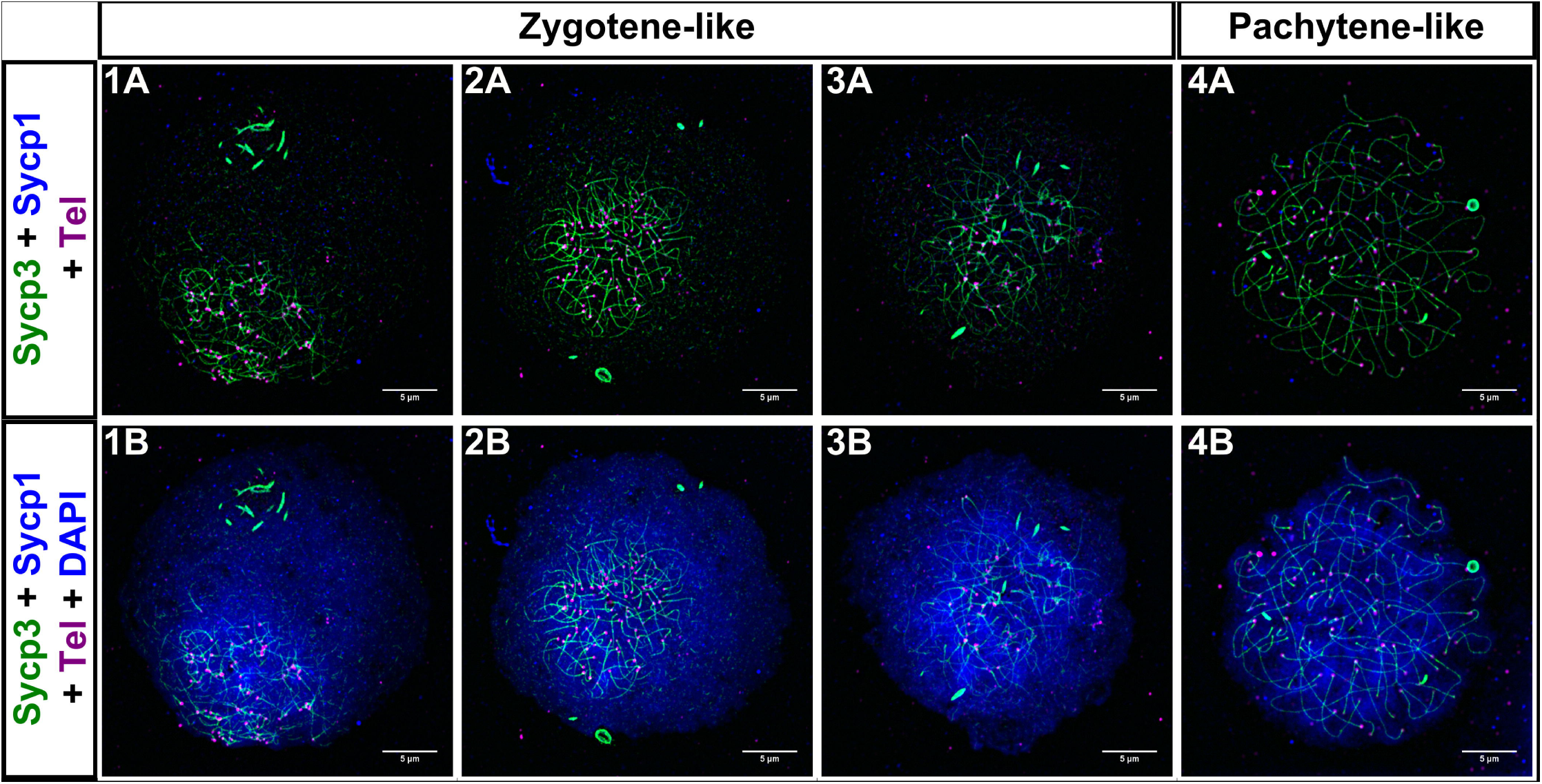
*spo11 -/-* females fail to properly synapse homologous chromosomes. 1A-4B: *spo11 -/-* nuclear surface spreads stained for Sycp3 (green), Sycp1 (blue), telomeres (Tel; magenta), and DNA (DAPI, blue). While Sycp3 loading proceeds, chromosomes are not synapsed. Scale bar = 5μm.)

### The *spo11* mutant males and females have sexually dimorphic reproductive defects

We found that *spo11* mutant males could induce spawning in females but failed to fertilize eggs (Sup. Fig. 4), indicating that they were either unable to produce or release their sperm. We inspected *spo11* mutant testes using light microscopy and found they appeared more translucent compared to wild type (Fig. 10A, panels 1A-2A), a phenotype that suggested a defect in sperm production. To confirm this, we isolated and stained whole testes with an antibody to the Vasa protein (Fig. 10A, panels 3A-4C). In wild-type zebrafish, Vasa is highly expressed in early germ cell clusters but diminishes as the spermatocytes progress in maturity, and is absent in mature spermatozoa clusters which can be identified by their tightly compacted nuclei [69]. We found that the *spo11* mutant males lacked sperm, and correspondingly, Vasa was expressed in all cell clusters, though it did diminish compared to the early germ cells.

**Fig. 10.**
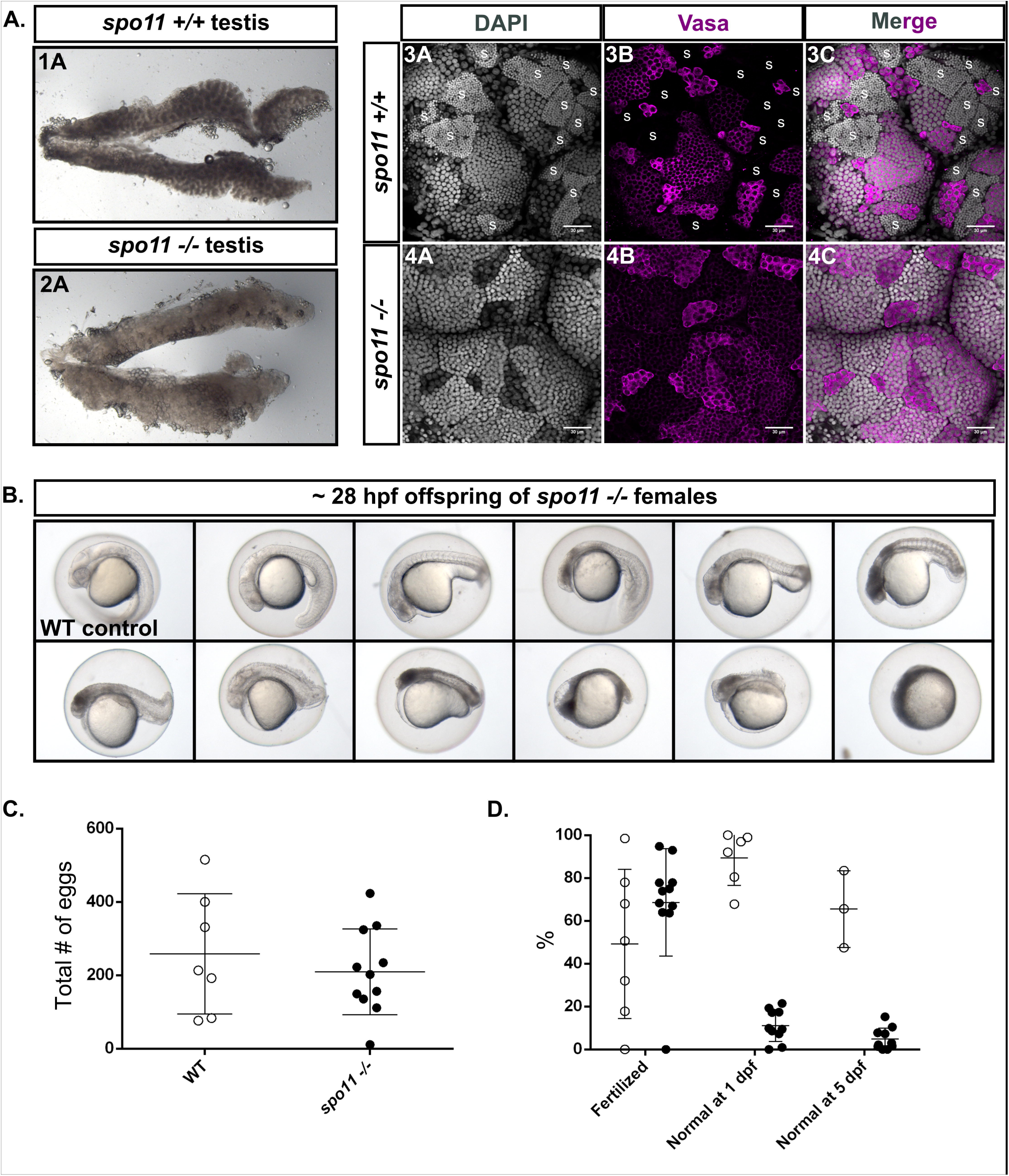
*spo11 -/-* males do not produce sperm, while *spo11 -/-* females produce offspring that show a spectrum of abnormalities. A: 1A-2A: Light microscope images of whole testes with *spo11 -/-* males showing a more translucent testis appearance as compared to WT. 3A-4C: Whole mount testes from WT and *spo11 -/-* males stained for Vasa (magenta) and DNA (DAPI, gray). Sperm clusters, indicated with “S” in WT, are absent in *spo11 -/-* mutants. B: Examples of offspring from *spo11 -/-* females showing the range of abnormalities at 28 hpf. hpf = hours post fertilization. C-D: Test crosses of *spo11 -/-* females (black circles) and WT control females (white circles) with WT males. Each circle represents a single test cross outcome. The data was pooled from two separate test crosses done on different days. The WT data at 5 dpf was assessed from one of the test crosses but was not tracked up to 5 days for the other WT test cross. C: *spo11 -/-* females produced similar numbers of eggs to WT females. D: The percent of eggs that were fertilized, and the percent of offspring that were normal or near normal in their development at 1 dpf, and at 5dpf. dpf = days post fertilization.

Surprisingly, *spo11* mutant females produced similar numbers of fertile eggs as wild type, however, the vast majority of their embryos died before 5 days post fertilization (dpf) and displayed a spectrum of abnormalities (Fig. 10B, C, and D). We expect that the severe developmental defects displayed among the progeny of *spo11* mutant females were a result of aneuploidy since it is unlikely that the chromosomes would be able to segregate properly with gross synapsis and pairing defects. The offspring that were normal at 5 dpf continued to grow into adults that developed as males. A similar offspring phenotype was seen in *mlh1* mutants in the zebrafish, where the offspring were shown to be aneuploid, and the ones surviving to adulthood developed as males that were found to be triploid [49]. Together, our data show that despite similar synapsis and pairing defects, males and females display dramatically different reproductive outcomes. This suggests a difference in checkpoint response between the sexes in the zebrafish.

## Discussion

### The bouquet is a hub for early meiotic chromosome events

Super-resolution analysis of homologous chromosome synapsis and pairing in the zebrafish revealed a coherent timeline of events (Fig. 11). 1) Assembly of the chromosome axis protein, Sycp3, initiates almost exclusively at both ends of chromosomes and elongates inward. 2) DSBs cluster near the telomere region. 3) Co-alignments form between telomere-proximal chromosome axes in funnel or pinch configurations. 4) The synaptonemal complex protein, Sycp1, loads between peri-telomeric axes and elongates slightly behind Sycp3 assembly. 5) Stable homolog juxtaposition at interstitial loci is not evident until the synaptonemal complex spreads across the region. As meiosis progresses, interlocks between chromosomes can be observed. Throughout the leptotene to pachytene stages telomere associations are present.

**Fig. 11.**
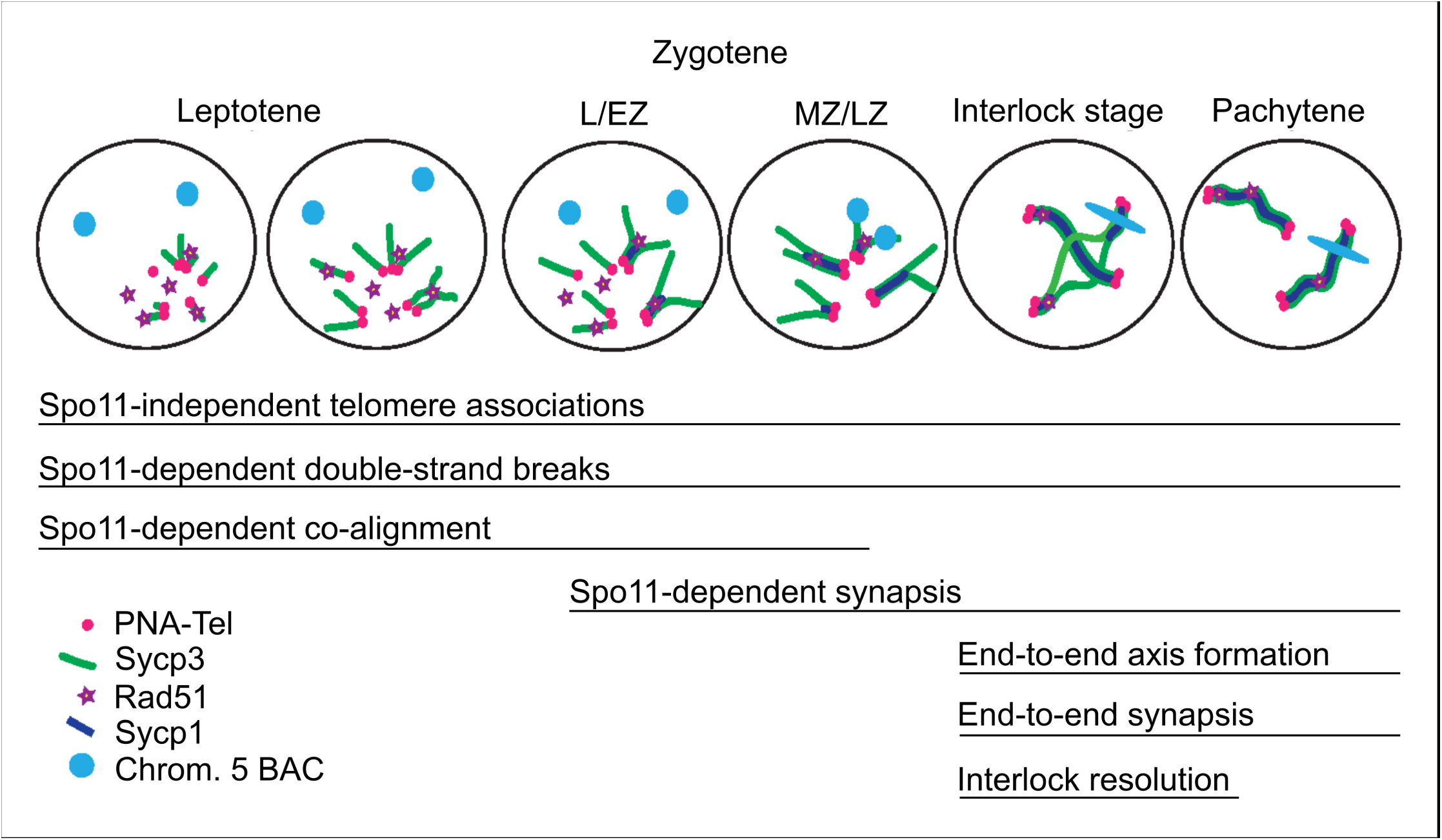
Model of the temporal progression of the chromosome events of meiotic prophase I. At leptotene, Sycp3 lines originate at the telomeres and elongate by extending toward the center of the chromosome. Initiation of Sycp3 growth is contemporaneous with the presence of Rad51 foci. Neither homolog alignment at interstitial regions nor synapsis are apparent, but some co-alignment of axes (less than 0.5 μm) near telomeres occurs. Zygotene stages, L/EZ and MZ/LZ, are defined in Figure 2 as leptotene/early zygotene and mid to late zygotene, respectively. At these stages, synapsis initiates between Sycp3 axes, yet not all Sycp1 lines are associated with a Rad51 focus. Close, stable homolog juxtaposition occurs as synapsis elongates from the telomeres. At a pre-pachytene stage, interlocks and/or regional de-synapsis occurs. As homologs are paired, the BAC signal becomes elongated, suggesting it is contained in a chromosome loop.

One of the most striking findings of our analysis was that the key events of meiotic chromosome metabolism, including axis morphogenesis, DSB formation, stable homolog juxtaposition, and synapsis all occurred within the limited region of the nucleus defined by the bouquet. Moreover, the focus of these events was specifically limited to the ends of chromosomes. From a first approximation, the general lack of close, stable homolog juxtaposition at interstitial sites suggests that the two ends of the same chromosome are not distinguished as such. Since zebrafish have 25 pairs of chromosomes, any given end is thus challenged to find its homologous partner among 99 possible choices within the bouquet prior to zygotene. Processes that promote the efficiency of pairing could include time intervals that favor collisions by diffusion [70, 71], rapid prophase movement to increase the rate of collisions via attachment of telomeres to cytoskeletal motor proteins outside the nucleus [36, 40, 72, 73] and/or through one or more DSB-independent pairing interactions [2, 27].

For organisms that initiate synapsis at sites of DSBs, homolog juxtaposition along the lengths of chromosomes can often be detected prior to synapsis (e.g. *Sordaria)[2]*. However, in these organisms a dramatic polarization of DSBs toward the telomere region, like that seen in zebrafish, is not evident. By contrast, DSBs and synapsis initiation in human males and the planarian *Schmidtea mediterranea* show a polarization similar to zebrafish [13, 74]. In the planarian, synapsis was shown to drive homolog pairing. It is possible that zebrafish homologs are paired by synapsis as well. A notable difference between the planarian and zebrafish, however, is that the planarian does not require Spo11 for synapsis whereas the zebrafish does, suggesting distinct SC nucleation methods between the two species. Our study does not answer the question of whether interstitial regions are physically juxtaposed by “zippering up” as SC spreads, or if a wave of DSBs creates new synapsis initiations sites and/or stabilizes SC.

Interestingly, in zebrafish, the bouquet is also the organizing center of the Balbiani body (Bb), a collection of embryonic patterning factors, mitochondria, and organelles which defines the animal-vegetal axis of the oocyte and is found in a wide variety of organisms including *Drosophila*, *Xenopus*, and mouse [75, 76]. In zebrafish, disruption of the bouquet *ex vivo* by the addition of the microtubule inhibitor nocodazole also disrupts Bb precursors showing the two structures are mechanistically linked [77]. It will be interesting to test if other meiotic chromosome features are also linked to the Bb, or if the Bb contributes to meiotic progression.

### Zebrafish offers new frontiers for studying regulation of chromosome dynamics

Our work uncovered several features of zebrafish biology that can stimulate new lines of enquiry to understand meiotic chromosome dynamics. First, nonhomologous telomere associations were prominent throughout meiosis, yet the nature of these associations is not well understood. One possibility is that they represent associations between heterochromatic regions like those seen in crickets [45], or telomere-bound protein interactions, as has been proposed for Trf1 [46]. We also do not know if they represent interactions between the same chromosomes from cell to cell. Associations could represent an early phase of the pairing process where the bouquet facilitates interactions between all telomeres, and rapid chromosome movements act to disrupt weak nonhomologous interactions to favor stronger DSB-dependent homologous interactions [32, 33, 36]. In addition, it is unknown what structure at or near the telomere seeds the initial loading of Sycp3, or the significance of the Sycp3 “bridges” sometimes seen between nonhomologous telomeres.

Second, our results show that Spo11 is required for the co-alignment of axes and SC formation. We attribute this effect to the formation of DSBs by Spo11 since the earliest occurrences of Rad51 and γH2AX signals are skewed toward telomeres where co-alignment and SC first appear. Observing Rad51 foci and γH2AX staining first near telomeres and later at interstitial locations suggests that DSBs may form in a wave, where initial breaks near telomeres bring homologs together to initiate local synapsis, while subsequent breaks form as Sycp3 is progressively loaded to initiate and/or to stabilize SC elongation. Consistent with the latter model, we occasionally see more than one synapsis initiation site between two chromosomes, albeit close to the telomere. A previous study showed that RPA foci, known to mark intermediates of DNA replication and DSB repair, form lines along the elongating axis in the zebrafish [56]. It is not known, however, if these are a result of DNA replication or Spo11-dependent recombination intermediates. Another possibility is that one or a few DSBs near the telomere are sufficient to promote synapsis along the length of a chromosome. The kinetic relationship between Sycp3 loading and Spo11-dependent SC initiation and elongation points to a possible regulatory mechanism that couples these processes. A study in the medaka fish, *Oryzias latipes*, has shown that loading of Sycp3 and Sycp1 is polarized to one side of the leptotene nucleus, with Sycp3 lines appearing to slightly anticipate Sycp1 [78]. This suggests that the mechanism of Sycp3 loading and synapsis in zebrafish may be common to other fishes.

Third, a transient interlock stage suggests a robust resolution mechanism. Interestingly, in cells where interlocks are observed, we also see pairs of chromosomes separated by long stretches of de-synapsed regions, sometimes disjoining two chromosomes completely. One possibility is that zebrafish employ long-range chromosome de-synapsis to resolve chromosome interlocks, as has previously been suggested in other organisms [62, 66]. It is possible that late-forming interstitial DSBs may play a role in re-establishing homologous synapsis at pachytene following this method of interlock resolution. In budding yeast, mouse, and *C. elegans*, SC components are involved in downregulation of DSB formation [7], thus it seems possible that local de-synapsis could activate new DSB formation.

Fourth, our analysis shows that synapsis initiates near the telomeres and progresses inward in both males and females, despite the differences in Mlh1 distribution and the recombination landscape between the two sexes [57, 58]. In mammals, the SC nucleation and crossover distribution landscapes correlate and are sex-specific (mouse, human, cattle) [20, 27–30]. In zebrafish, it is possible that the relationship between SC nucleation and crossover designation differs between the sexes, given that in males the SC nucleation pattern resembles the crossover pattern whereas in the females it does not appear to.

### Zebrafish as a model for sex-specific reproductive defects and checkpoint function

Zebrafish is not only an excellent model to study the events of meiosis *per se*, but also to study sexually dimorphic responses to meiotic perturbations. Zebrafish has previously been proposed as a model for germ cell aneuploidy [50, 79]. We show here that despite exhibiting similar defects in synapsis and pairing, *spo11* mutant males and females show vastly different outcomes in reproduction. The males are unable to produce sperm, while females produce eggs that result in severely deformed offspring. This is in line with previous studies that show sexually dimorphic outcomes: Disruptions of Mlh1 [49, 51], and Mps1, a kinase required for the spindle assembly checkpoint [50], show a tendency for females to produce aneuploid offspring. Unlike in *spo11* mutants, where males are sterile, and *mlh1* mutants, where males are predominantly sterile, both male and female *mps1* mutants produce aneuploid offspring, although the rate is higher in females than males (~46% vs. ~26% respectively). This suggests complex mechanisms underlying causes of increased aneuploidy in zebrafish females.

Sex specific differences are also seen in *spo11* mutant mice; spermatocytes die by early pachytene whereas oocytes survive until diplotene/dictyate stage [80, 81]. The arrest seen in mouse males, however, is likely different than the arrest seen in zebrafish. Among organisms with heterogametic sex determination, mechanisms have evolved to specifically accommodate unpaired chromosomes in the heterogametic sex, including meiotic sex chromosome inactivation [82, 83]. As such, mutations that disrupt pairing might be expected to have a weaker effect in the homogametic sex, where the MSCI checkpoint may not be as robust [84]. In domesticated zebrafish, sex determination is polygenic, with no universal structural differences between chromosome sets of sexes in lab strains [58, 85]. Consistent with these findings, we did not observe any chromosomal regions that remained unpaired during pachytene. Thus, the pronounced effect of the *spo11* mutation in males is likely not due to the activation of the MSCI checkpoint. Instead, the failure to produce sperm may depend on another checkpoint, such as the synapsis or the spindle assembly checkpoints, that operate in other model systems [1, 86–89].

### Zebrafish as a model for human reproduction

Our findings highlight the importance of studying multiple model systems. While homolog pairing and recombination are considered universal features of meiosis, the means to getting there is quite varied among species. Interestingly, some meiotic prophase events in zebrafish resemble the corresponding events in human spermatogenesis, including the tendency of DSBs to skew near the ends of chromosomes and the initiation of synapsis at telomeres followed by inward synaptic progression [74, 90]. Telomere-proximal synapsis initiation while Sycp3 loading is not yet complete has also been reported in human spermatocytes [19, 20]. However, in humans the Sycp3 loading appears more extensive than in the fish by the time that synapsis ensues. Understanding spermatogenesis is important since sperm concentration and total sperm count has declined 50–60% between 1973 and 2011 among men in western countries [91], and the causes behind male infertility remain unknown in about 40% of patients [92]. In addition, human females are more prone to generating aneuploidy as compared to males [1, 93–95], which resembles the situation in zebrafish. While the causes of aneuploidy and reduced fertility in humans are complex and the contributions are manifold, zebrafish could provide valuable insights into environmental, genetic, and sex-specific effects on adverse meiotic outcomes.

## Methods

### Zebrafish Strains

The wild type AB strain was used in the production of *spo11* mutants. Wild type data presented in figures 1 - 6 and 10 - 11 are from tank mates of *spo11* mutants. AB strain fish were used for figure 7A, and NHGRI strain fish were used for figure 7B. NHGRI strain fish were used for test crosses in one of the two pooled data sets in figure 11. Other test crosses were done with AB strain fish. Fish were maintained as previously described [96].

### Mutant generation and identification

Spo11 mutants were generated using TALENs targeting the second exon of *spo11*. The TALENs were assembled and injected as previously described [97]. The TALEN sequences were: HD-NG-NI-NI-NI-NN-NN-NG-NN-NI-NI-NN-HD-NI-HD-half repeat HD, and NG-HD-HD-NI-NN-HD-NI-NN-NN-NI-NG-HD-NG-NI-NG-half repeat NG. Injected founder fish were raised to adulthood and outcrossed to wild type fish; the resulting offspring were screened for mutations in *spo11* via high resolution melt (HRM) analysis and subsequent sequencing. HRM primer sequences are: fwd TCACAGCCAGGATGTTTTGA, and rev CACCTGACATTGTTCCAGCA. The HRM analysis was performed with either Light Scanner Master Mix (BioFire Defence, Murray, UT, Catalog# HRLS-ASY-0003), 10X LCGreen Plus+ Melting Dye (Biofire Defence, Catalog# BCHM-ASY-0005), or 20X Eva Green dye (VWR, Radnor, PA, Catalog# 89138–982) using a CFX-96 real-time PCR machine and Precision Melt Analysis software (BioRad, Hercules, CA). The data presented in this paper is from individuals of a population with an 11 bp deletion mutation in exon 2 that has been outcrossed 2 - 3 times.

### Reproducibility

All our conclusions are based on experiments that were performed at least two times. All data sets comparing WT and *spo11* mutants were collected from tank mates processed in parallel on the same days, including the spreads and staining the slides. The antibodies, the BAC probe and the Telomere PNA probes were tested multiple times on spreads and/or whole-mount gonads prepared on different days. The Student t-test was used for statistical analysis. All numerical data used for each plot is tabulated in Supplemental table 1. Raw SIM data for all cells are available upon request. Images shown in each figure will be deposited at the Dryad Digital Repository (https://datadryad.org/).

### Adult testes chromosome spreads

About 15 - 20 gonads were freshly dissected in 1X Phosphate Buffered Saline (PBS). The gonads were placed in 2 ml Dulbecco’s Modified Eagle Medium (DMEM) in a 5 ml Eppendorf tube on ice. 4 mg of collagenase (Sigma-Aldrich Chemical Co Inc, St. Louis, MO, Catalog# C0130–500MG) dissolved in 200 μl DMEM were added and the gonads were gently shaken horizontally at 32°C for 50 minutes to an hour, until the liquid was cloudy, and the gonads were in small chunks. The tube was inverted rapidly several times every 10 minutes to facilitate dissociation. The collagenase was then washed out: DMEM was added up to 5 ml and the gonads were pelleted at 218g for 3 minutes. Then 3 ml of the supernatant were removed to reduce the liquid down to 2 ml. This was repeated 2 additional times for a total of 3 DMEM washes with the supernatant reduced to 1 ml after the last wash (the pellet was not resuspended between the washes).

DMEM was added up to 2 ml total, and 1.4 mg trypsin (Worthington Biochemical Corporation, Lakewood, NJ, Catalog# LS003708) dissolved in 200 μl DMEM and 20 μl of 400 μg/ml DNaseI (Roche Diagnostics, Pleasanton, CA, Catalog# 10104159001) were added for cell dissociation. The tube was gently shaken horizontally at 32°C for 5 - 15 minutes until the solution contained few clumps. The tube was inverted rapidly several times every 5 minutes to facilitate dissociation. 10 mg of trypsin inhibitor powder (VWR, Catalog# IC100612.5) dissolved in 500 μl DMEM and 50 μl of 400 μg/ml DNase I solution were then added. The tube was briefly spun down, and the cell suspension was pipetted repeatedly up and down with Pasteur pipettes for 2 minutes to facilitate dissociation of any remaining clumps.

The cell suspension was put through a 100 μm nylon Falcon filter (Fisher Scientific, Waltham, MA, Catalog# 08–771–19) and transferred to a fresh 5 ml tube. DMEM was added to 5 ml total volume and the cells were pelleted at 218g for 5 minutes. The supernatant was removed and 5 μl of the DNase I solution was added directly to the pellet which was then resuspended by scraping the bottom of the tube on an empty tube rack. DMEM was added up to 5 ml and the cells were pelleted at 218g for 2 minutes. The DNase I treatment was repeated a total of 2 - 4 times until the resuspended pellet did not clump upon addition of DMEM. After the last treatment, the pellet was resuspended in 1 - 2 ml of 1X PBS and pelleted again at 218g for 5 minutes. The supernatant was removed and the pellet resuspended with a pipette tip (~3 mm cut off from tip to widen the aperture) in 80 - 100 μl of 37°C 0.1M pH ~8 sucrose, and allowed to sit at room temperature for 3 - 5 minutes.

Slides (Fisher Scientific Premium Superfrost, Catalog# 12–544–7) were coated with 100 μl of 1% Paraformaldehyde (PFA; Acros Organics, Catalog# 30525–89–4) with 0.15% Triton X-100 (Fisher BioReagents, Catalog# 9002–93–1) and then ~20 μl of cell suspension was added directly to the center of the slide in a straight line. The slide was tilted to facilitate spreading. The slides were placed in a slightly cracked open flat humid chamber. The chamber was placed in a dark drawer and allowed to sit overnight. It was then opened, and the slides allowed to completely dry. The slides were rinsed for 5 minutes in H_2_O and then twice for 5 minutes in 1:250 Photo-Flo 200 (Electron Microscopy Sciences, Catalog# 74257) in Coplin jars. The slides were dried and stored at −20°C until they were stained.

### Adult ovary chromosome spreads

About 6 - 10 gonads of females aged 60 - 80 dpf were dissected in 1X PBS. The gonads were placed in 2 ml DMEM in a 5-ml tube and passed through an 18-gauge needle and then a 20-gauge needle 15 times each. The cells were briefly spun down and then pipetted up and down with Pasteur pipettes for 2 minutes. The cell suspension was put through a 100 μm nylon Falcon filter and transferred to a clean 5 ml tube. The cells were then pelleted at 218g for 5 min. The pellet was composed of two layers, a bottom whitish layer and a top yellowish layer. The top layer was carefully removed with pipette and the remaining bottom layer was resuspended in 2 ml 1X PBS. The cells were then pelleted at 218g for 5 min, the supernatant was removed, and the pellet was resuspended with cut pipette tip in 80 - 100 μl 37°C 0.1M pH ~8 sucrose. The suspension was allowed to sit at room temperature for 3 - 5 minutes. The slides were prepared as in the *“Adult testes chromosome spreads”* protocol.

### Telomere probe and hybridization solution preparation

PNA telomere probes TelC-Alexa647 and TelC-Cy3 were acquired from PNA Bio Inc, Thousand Oaks, CA (Catalog# F1013 and F1002 respectively); 50 μM stocks were prepared in formamide as per manufacturer’s instructions and stored at −80°C. The hybridization solution was prepared to a final concentration of 0.2 μM PNA telomere probe and 1.33 mg/ml bovine serum albumin (Fisher Scientific) in pre-hybridization solution. The pre-hybridization solution was composed of 50% formamide (Fisher Scientific, Catalog# BP228–100), 5X Saline-Sodium Citrate (SSC; 20X stock: 3M NaCl and 0.3M Sodium Citrate), 50 μg/ml Heparin sodium salt from porcine intestinal mucosa (Sigma-Aldrich Chemical Co Inc, Catalog# H3393–100KU), 500 μg/ml transfer RNA from wheat germ (Sigma-Aldrich Chemical Co Inc, Catalog# R7876–2.5KU), 0.1% Tween 20 (Bio-rad, Catalog# 170–6531), and 1M Citric acid to bring the solution to pH ~6. The pre-hybridization and hybridization solutions were stored in −20°C in the dark.

### Chromosome spreads staining

#### PNA telomere probe staining

Chromosome spread slides were re-hydrated for 5 minutes in ~500 μl 1X PBS which was removed afterwards by tapping the side of the slide on a paper towel. The slides and hybridization solution were preheated separately to ~82°C - 83°C in a hybridization oven. 100 μl of hybridization solution was then added per slide and covered with a plastic coverslip (cut to shape out of an autoclave bag). The slides were heated at ~82°C - 83°C for 10 - 12 minutes, and then placed in a flat humid chamber in the dark at 37°C overnight (~16 - 24 hours). The coverslips were removed, and the slides were washed for a minimum of 15 minutes per wash in 1) pre-hybridization solution with no transfer RNA added covering the slide, 2) 50% pre-hybridization solution with no transfer RNA added in 1X PBS covering the slide, and 3) 3 times in 1X PBS with gentle shaking in a coplin jar. The slides were taken out of the jar and tapped on a paper towel to remove excess PBS.

#### Primary antibody staining

The slides were blocked in a humid chamber at room temperature for a minimum of 20 min with the antibody block covering the slide. Antibody block was prepared to a final concentration of 2 mg/ml BSA and 2% Normal goat serum (NGS; Sigma-Aldrich Chemical Co Inc, Catalog# G9023–10ML) in PBT (1X PBS + 0.1% Triton X-100). The block was removed and 100 μl of primary antibody mix in antibody block was added per slide and covered with a plastic coverslip. The primary antibodies and their working concentrations were: 1:200 Rabbit anti-SCP3 (Abcam, Cambridge, United Kingdom, Catalog# ab150292), 1:100 Chicken anti-Sycp1 (see methods below), and 1:50 Rabbit anti-Rad51 (AnaSpec, Fremont, CA, Catalog# old: 55838–2, new: AS- 55838). The slides were placed in a dark, humid chamber overnight at 4°C. The coverslips were removed, and the slides were washed 2 times in 1X PBS in a gently shaking coplin jar for a minimum of 5 minutes each.

#### Secondary antibody staining

Excess PBS was removed, and the slides were blocked at room temperature for a minimum of 5 minutes with the antibody block covering the slide. The block was removed and 100 μl of secondary antibody mix in antibody block was added per slide and covered with a plastic cover slip. The slides were placed in a flat, humid chamber at 37°C for 1 hour. All secondary antibodies were added at 1:1000 final concentration. The following secondary antibodies were acquired from Thermo Fisher Scientific Inc: Goat anti-Rabbit IgG (H+L) Cross-Adsorbed Secondary Antibody, Alexa Fluor 488 (Catalog# A-11008), Goat anti-Rabbit IgG (H+L) Cross-Adsorbed Secondary Antibody, Alexa Fluor 594 (Catalog# A-11012), Goat anti-Chicken IgY (H+L) Secondary Antibody, Alexa Fluor 488 (Catalog# A-11039), and Goat anti-Chicken IgY (H+L) Secondary Antibody, Alexa Fluor 594 (Catalog# A-11042). The following secondary antibody was obtained from Biotium Inc, Fremont, CA: CF®405M Goat Anti-Chicken IgY (H+L), highly cross-adsorbed (Catalog# 20375–500uL). The coverslips were removed, and the slides were washed 3 times in 1X PBS for a minimum of 5 minutes each. They were then washed for a minimum of 2 minutes in H2O and allowed to dry completely in the dark, tilted to facilitate drying. The slides were then mounted with 20 μl of Thermo Fisher Scientific ProLong™ Diamond Antifade Mountant with DAPI (Catalog# P36966), or without DAPI (Catalog# P36965). The prepared slides were stored at 4°C.

### Bacterial Artificial Chromosome (BAC) Probe DNA Preparation

BAC clone CH211–31P3 (https://zfin.org/ZDB-BAC-050218–850) was obtained from the BACPAC Resources Center (BPRC). The BAC was purified via Midiprep as previously described [98]. Purified BAC quality was assessed by running a sample on a 1% agarose gel, and the BAC’s identity was confirmed by PCR amplification of a segment of the *nanos2* gene using the following primers: fwd ATGCAGTCCGAGAGTCAGCAGAG, and rev ATAACGGACACACGTAGCTCCTCAG.

### Cot-1 preparation

The Cot-1 preparation was adapted from [98]. Salmon testes DNA (Sigma-Aldrich, Catalog# D1626–1G) was prepared at 10 mg/ml in H_2_O by dissolving overnight at 55°C. 300 μl aliquots of the testes DNA were sonicated in Diagenode tubes (Fisher Scientific, Catalog# NC0065146) in a Diagenode Bioruptor UCD-300 for ~60 15-second cycles or until the average fragment size was ~400–500 bp. The fragment size was checked on a 1% agarose gel. 500 μl of sonicated salmon testes DNA was denatured at ~100°C for 15 minutes and then incubated at 65°C for 4 minutes. 250 μl of 1M NaCl (pre-heated to 65°C) was added and the mix was incubated at 65°C for the duration of time needed for the Cot-1 fraction to re-anneal (equation: 5.92/DNA concentration in mg/ml = time (in minutes)). Then 1 unit of S1 nuclease (Thermo Fisher Scientific, Catalog# EN0321) per 1 μg of DNA was added together with 5X S1 nuclease reaction buffer. The mixture was incubated at 37°C for 30 minutes.

The Cot-1 solution was transferred to a 15 ml conical tube, mixed with 10 ml of pH 8 Phenol:Chloroform:Isoamyl alcohol 25:24:1 (Fisher Scientific, Catalog# BP1752–100) and centrifuged at 1500g for 5 minutes. The aqueous phase was transferred to a new 15 ml tube and mixed thoroughly with 0.1X volume of 3M sodium acetate. 1X volume of 100% isopropanol was added, the solution mixed gently to precipitate the DNA, and then centrifuged at 3000g for 10 minutes at 4°C. The supernatant was removed, and the pellet was allowed to air dry with the tube inverted at an angle. The pellet was re-hydrated in 30 μl H_2_O and the concentration was determined by nanodrop. The pellet was further cleaned with 1 ml Phenol:Chloroform:Isoamyl alcohol 25:24:1 followed by 70% EtOH. The final pellet was dried and resuspended in 30 μl H_2_O and the concentration was determined by nanodrop.

### BAC probe labeling and preparation

The BAC probe labeling and preparation was adapted from [98]. The probe was labelled with Green dUTP (Abbott Molecular, Abbott Park, IL, Catalog# 02N32–050) using the Nick Translation Kit (Abbott Molecular, Catalog# 07J00–001). 14 μl of the purified BAC was mixed with 23.4 μl of 0.1 mM dNTP mix (1:2:2:2 of dTTP:dATP:dCTP: dGTP), 10 μl of 10X Nick translation buffer, 10 μl of the Nick translation enzyme mix, 12 μl of 0.2 mM Green dUTP, and H_2_O to bring up the volume to 100 μl. The reaction was incubated in a thermocycler at 15°C for 16 hours, heated to 70°C for 10 minutes, and then held at 4°C. The labeled BAC was purified using DNA Clean & Concentrator™-5 (Zymo Research, Irvine, CA, Catalog# D4013) in 50 μl batches and eluted in 10 μl of the elution buffer. 25 μg of salmon sperm Cot-1 was added per batch and the batches were mixed together. The mixture was vacuum dried, and the pellet was resuspended in 10 μl of LSI buffer (LSI/WCP^®^ Hybridization Buffer, Abbott Molecular, Catalog# 06J67–011) to make the stock BAC probe mix. The stock was stored in the dark at −20°C. For staining, the stock was further diluted in LSI buffer at a 1:19 stock:LSI ratio.

### BAC probe staining

The BAC probe staining procedure was adapted from [98]. Chromosome spread slides were placed in 3:1 MeOH:HAc at −20°C for 15 minutes. The slides were then washed 2 times in 1X PBS for a minimum of 2 minutes each and treated with 0.5 mg/ml Protease II (Abbott Molecular Inc., Catalog# 06J93–001) at 37°C for 5 minutes. The slides were washed 2 times in 1X PBS for a minimum of 5 minutes each, and then progressively dehydrated in 2-minute washes with 70%, 85%, and 100% EtOH. The slides were allowed to air dry completely and used immediately for staining.

Prior to BAC probe staining, PNA telomere probe staining was performed as described in the “*PNA telomere probe staining*’ section, and the slides were allowed to air dry completely after the final 1X PBS wash. At this point, 10 μl of the BAC probe (1:19 dilution in LSI buffer) was added per slide, covered with a 24 × 50 coverslip, and sealed with rubber cement (Elmer’s, Atlanta, GA, Catalog# E904). The slides were heated in a hybridization oven at ~70 - 71°C for 3 minutes and then the oven temperature was allowed to drop to ~50°C after which the slides were transferred to a flat, humid chamber and incubated at 37°C overnight in the dark. The coverslip was peeled off and the slides were washed in coplin jars in 1) 50% formamide in 2X SSC at 45°C, 2 times for 5 minutes each, 2) 2X SSC at 45°C, 2 times for 5 minutes each, 3) 4X SSC + 0.05% Tween 20 for 8 minutes, 4) 1:1 2X SSC:PBSTw (1X PBS + 0.1% Tween 20) at room temperature (RT) for 5 minutes, and 5) PBSTw at RT, 3 times for 5 minutes. Excess PBSTw was removed from the slides by tapping their sides on a paper towel. The antibody staining and slide mounting were performed as described in the *“Primary antibody staining*” and “*Secondary antibody staining*” sections, with PBSTw used instead of PBT.

### Chromosome spreads measurements

#### Synaptonemal complex/ staging

Sycp3 lines were measured from the telomere to the end of the Sycp3 stained region using the segment tool of ImageJ. Only continuous Sycp3 lines with no significant gaps in signal were measured. Similar analysis was done for Sycp1 lines. In the limited number of cases where an Sycp1 line originated not immediately adjacent to the telomere PNA probe, these distances were included in the measurements.

#### BAC probe distance

The segment tool of imageJ was used to draw an outline of each BAC signal to find the (x,y) coordinates of the centers of mass. The distance between the two points was calculated using the distance formula 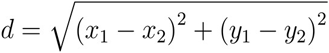.

#### Co-alignment

The co-alignments were measured according to the following criteria:

1. Telomere proximal Sycp3 labeled axes appear to be coming together in preparation for synapsis.
2. The maximum distance between the co-aligning axes at the narrowest region is 0.5 μm (other organisms show co-alignment at 0.4 μm, an extra 0.1 μm was added to allow for human error in measurement).
3. We classified the co-alignment shape types into two broad categories: funnel and pinch. In the funnel configuration Sycp3 lines have a co-aligned region directly adjacent to the telomeres and the Sycp3 lines extend past the co-aligned region forming a funnel-like structure; the Sycp3 lines past the co-alignment can be bent in any direction or can cross each other. In the pinch configuration the narrowest region between the Sycp3 axes is slightly away from the telomeres forming a pinch-like structure.
4. In cases of crowded areas in the chromosome spread it is clear which axis belongs to which telomere; both telomeres can be identified in the structure even if one is much dimmer than the other.
5. Slightly misaligned telomeres in otherwise co-aligned axes are permitted since the orientation of the chromosomes may shift when the 3D nucleus is flattened to 2D.
6. Sycp3 labeled axes that have a region of Sycp1 connecting the two Sycp3 axes together anywhere between them were not considered co-aligned as they have already initiated synapsis. As the Sycp1 antibody may have varying amounts of background, we assessed the signal between the Sycp3 lines by applying the MaxEntropy Auto Threshold method in ImageJ to the Sycp1 channel to standardize our assessment between different images.

#### Telomere associations

The telomere associations were measured as numbers of engaged telomeres using the following criteria:

One engaged telomere end consisted of a telomere signal + Sycp3 line.
1. Telomeres could be round or elongated.
2. Telomeres were either touching or connected by an Sycp3 bridge.
3. If telomeres of synapsed chromosomes were engaging with a non-homolog, only 1 end of the synapsed chromosome was counted as associated, as the other end may have been there due to synapsis with the first end and not an association with the non-homolog.
4. If the interaction between telomere ends looked like a co-alignment or if there was any synapsis between the Sycp3 lines, the telomere ends were not counted as associated.
5. If the telomeres were touching but the sycp3 lines were crossing directly behind the telomeres they were not counted as associated since that could have been a co-alignment in another plane view.

### Whole mount testes staining

#### Dissection and fixation

Whole mount testes antibody staining was adapted from [99]. Euthanized fish were decapitated and cut open along the ventral midline to expose the body cavity. The fish were fixed in 4% PFA in PBT at 4°C overnight (~16 - 18 hours) with gentle rocking. The fish were washed in PBT 2 times for ~10 minutes each and then the testes were dissected in PBT and placed in 1.5 ml centrifuge tubes. Alternatively, the testes were dissected in PBT prior to the 4% PFA fixation overnight, and then washed 2 times in PBT for ~10 minutes. Whether fish or dissected testes were fixed overnight did not affect subsequent staining.

#### Primary antibody staining

The testes were washed 3 - 4 more times for a minimum of 5 minutes each in PBT. Testes that were to be stained with the rabbit anti-γH2AX antibody [100] and chicken anti-Vasa (see methods below) were dehydrated in 2 washes of 100% MeOH for 5 minutes each, and then stored in 100% MeOH at −20°C for 1 hour minimum. The testes were rehydrated with a series of 5-minute MeOH washes with 75%, 50%, and 25% MeOH respectively, and then washed 5 times for 5 minutes minimum in PBT. The testes were then washed in antibody block (same recipe as for the “*chromosome spreads protocol*”) for 1 hour minimum with gentle rocking at room temperature. Testes that were stained with the rabbit anti-Vasa only [101], proceeded directly to the antibody block step. After blocking, primary antibodies were added at final dilutions of 1:100 for anti-γH2AX, and 1:500 for the anti-Vasa antibodies, and the tubes were rocked gently overnight at 4°C.

#### Secondary antibody staining

The testes were washed 4 times for a minimum of 15 minutes and 2 times for a minimum of 30 minutes in PBT, and then washed in antibody block as described above. Secondary antibodies, anti-rabbit Alexa Fluor 488 and anti-chicken Alexa Fluor 594 (same as for the “*chromosome spreads protocol*”) were added at final dilutions of 1:300 for the anti-γH2AX and the chicken anti-Vasa antibodies, and 1:500 for the rabbit anti-Vasa antibody and rocked gently overnight at 4°C.

#### Glycerol dehydration and mounting

The testes were washed the same as after the primary antibody staining and dehydrated in a series of glycerol (Sigma-Aldrich, Catalog #: G5516–1L) washes for 1 hour minimum each: 30% glycerol with DAPI in PBT or PBSTw, 50% glycerol with DAPI in PBT or PBSTw, and 70% glycerol in PBT or PBSTw with or without DAPI. The testes were mounted in 70% glycerol without DAPI on slides with vacuum grease applied to the four corners to hold the coverslip in place.

### Testes section preparation and staining

#### Testes section preparation

Fish were prepared as described in “*whole mount testes staining*” and fixed overnight in 4% PFA, and then washed 2 times for 15 min each in PBT. The testes were dissected and washed in PBT 2 times for 10 min each. The testes were dehydrated in a series of 1-hour minimum EtOH washes at 70%, 80%, 95% and 100% EtOH with 2 washes per concentration. The testes were then washed with 50:50 100% EtOH:Histoclear (National Diagnostics, Atlanta, GA, Catalog#: 5989–27–5) for 1 hour minimum, and then 2 times for 1 hour minimum in Histoclear. The testes were then washed 4 times for 1 hour each minimum in ~60°C paraffin wax (Paraplast plus, Electron Microscopy Sciences, Hatfield, PA, Catalog#: 19216), after which they were embedded in the paraffin wax and 5 μm sections were made using a microtome. The sections were heated in a water bath at 42°C to straighten out the wax strips, mounted on superfrost plus slides (Fisher Scientific, Catalog#: 12–550–15) and allowed to dry overnight. The sections were stored at 4°C until ready to stain.

#### Section staining

The sections were deparaffinized in 2 washes of histoclear for 5 mins each, followed by 2 100% EtOH washes for 3 mins each, a 1 min wash in 95% EtOH, and a rinse in water. The slides were then washed 3 times for 2 mins each in PBSTw. The slides and the PNA probe hybridization buffer were pre-heated to 85°C for 2 mins, 100 μl of the hybridization buffer was added to the slide, and the slide was covered with a plastic coverslip and incubated at 85°C for 10 mins. The slides were then incubated at 37°C overnight in the dark. The coverslips were removed, and the slides washed 3 times for 15 min each with 500 μl of pre-hybridization buffer without tRNA added to the top of the slides. The slides were then washed 2 times for 15 mins each in PBSTw or PBS, dried, mounted in Prolong Diamond Antifade and checked for telomere staining. The slides were then washed 2 times for 15 mins in PBT and PBSTw and blocked with antibody block for 1 hour minimum prior to antibody staining. 100 μl of the primary anti-γH2AX antibody was added at 1:100 concentration in antibody block and incubated overnight at 4°C in the dark with coverslip. The slides were then washed 3 times, 2 min each in PBSTw, and then blocked in antibody block for 1 hour minimum. ~150 μl of secondary antibody Alexa-fluor 488 was added at 1:300 concentration, covered with coverslip, and allowed to sit at room temperature for 1 hour in the dark. The slides were washed 3 times for 2 mins each, allowed to fully dry, mounted with Prolong Diamond Antifade with DAPI, covered with a glass coverslip, sealed with nail polish along the frosted edge of the slide, and stored at 4°C in the dark until ready for imaging.

### Imaging

All images were collected at the Department of Molecular and Cellular Biology Light Microscopy Imaging Facility at UC Davis. Chromosome spreads were imaged using the Nikon N-SIM Super-Resolution microscope in 3D-SIM imaging mode with Apo TIRF 100X oil lens. The images were collected and reconstructed using the NIS-Elements Imaging Software. Sections and fluorescent whole mounts were imaged using the Olympus FV1000 laser scanning confocal microscope. Images were processed using the Fiji ImageJ software. Only linear modifications to brightness and contrast of whole image were applied. All raw image files are available upon request.

### Test crosses

To analyze fertility, individual mutant fish were crossed to wild type fish to assess their ability to generate offspring. Offspring that were produced were tracked daily for up to 5 days to assess morbidity and mortality.

### Chicken anti-zebrafish Ddx4/Vasa polyclonal antibody production

A cDNA fragment of zebrafish ddx4/vasa encoding the COOH-terminal amino acids 479–651 (based on accession number BC129275) was cloned into pET100 using the following primers: fwd 5’-CACCATGTTCATAGCAACATTTCTCTGTCAAG-3’ (ATG initiation codon added); rev 5’-TAACAGGTGTGAGGCCAGTTATTCC-3’. The His-tagged protein was expressed in E. coli, purified using standard procedures and used to immunize chicken hens (91-day protocol, Pocono Rabbit Farm & Laboratory Inc. Canadensis, PA). Polyclonal IgY from crude serum was used at 1:500.

### Chicken anti-zebrafish Sycp1 polyclonal antibody production

An N-terminal fragment of Sycp1 cDNA was amplified with Phusion DNA polymerase (Thermo Fisher Scientific, Catalog#: M0530L) using the following primers: Fwd 5’-aactttaagaaggagatataccATGCAAAAAGCATTCAACTT-3’, and Rev 5’-tctcagtggtggtggtggtggtgctcGGTAACTTCTATTTCTGCATtt-3’. The Sycp1 PCR product (1272 bp) was then cloned into pET28b using NEBuilder HiFi DNA Assembly Master Mix (NEB, Ipswich, MA, Catalog#: E5520S). BL21 (DE3) cells containing pRARE and Sycp1 overexpression construct were grown in 2.6 L of LB with kanamycin and chloramphenicol until an OD600 = 1 and induced with a final concentration of 1 mM IPTG at room temperature for six hours. The Sycp1 peptide was purified under denaturing conditions using Novagen NiNTA purification resins (Sigma, Catalog#: 70666) according to the manufacturer’s instructions. The Sycp1 peptide was concentrated to a final concentration of 1mg/ml in PBS using a 10kDa centrifugal filter (Sigma, Catalog# UFC901008). The Sycp1 peptide was injected into two chickens by Pocono Rabbit Farm and Laboratory following the 91-day polyclonal antibody production protocol.

## Acknowledgements

We thank Rebecca Beer and Stephanie Lan-Uyen Nguyen for technical assistance with anti- Ddx4/Vasa antibody production; Daniel Chu for technical assistance with anti-Sycp1 antibody production; James Amatruda for the γH2AX antibody; Irina Vaysertreyger and Hester Roberts for assistance with the optimization of the BAC probe protocols; Prasada Rao for assistance with the optimization of the chromosome spreads protocol; and Roberto Pezza and Kelly Komachi for insightful comments on the manuscript. We thank Michael Paddy for advice and technical support at the MCB Light Microscopy Imaging Facility at UC Davis

---

**Sup. Fig. 1.** The Synaptonemal Complex (SC). A: A schematic representation of the SC. Both sister chromatids are tethered to one axis via cohesins. Sycp1 proteins form dimers and assemble into transverse filaments holding the two axes together. Sycp1 N-termini are in the middle and the C-termini are at the axes. A number of proteins are loaded at the center to form the central element of the SC. B: A representative SC of the zebrafish, *Danio rerio*. Chromosome loops, marked by DAPI (blue), are tethered to the axis, marked by an antibody to Sycp3 (green). The transverse filaments are marked by an antibody to the N-terminal regions of Sycp1 (magenta).

**Sup. Fig. 2.** BAC probe staining (red) shown separately with the telomere (Tel; magenta), Sycp3 (green), and Sycp1 (blue) channels. Same images as shown in Fig. 3. Scale bar = 5 μm.

**Sup. Fig. 3.** *spo11 -/-* male and female nuclear surface spreads stained for Sycp3 (green), Sycp1 (blue), telomeres (Tel; magenta), and a region ~10Mb from the telomere on chromosome 5 (BAC probe; red in merge channels, gray in separate channels). The BAC signals remain unpaired, indicating lack of homolog pairing. Scale bar = 5μm.

**Sup. Fig. 4.** *spo11* -/- males induce spawning but are unable to fertilize the eggs. Test crosses of *spo11* -/- males (black squares) and WT control males (white squares) with WT females. Dot plots show the number of eggs produced by females crossed to the test males (left), and the percent of the eggs that were fertilized (right). Each square represents the outcome for one male.

